# Shallow networks run deep: Peripheral preprocessing facilitates odor classification

**DOI:** 10.1101/2023.07.23.550211

**Authors:** Palka Puri, Shiuan-Tze Wu, Chih-Ying Su, Johnatan Aljadeff

**Author notes:** To whom correspondence should be addressed; (J.A.).

## Abstract

The mammalian brain implements sophisticated sensory processing algorithms along multilayered (‘deep’) neural-networks. Strategies that insects use to meet similar computational demands, while relying on smaller nervous systems with shallow architectures, remain elusive. Using *Drosophila* as a model, we uncover the algorithmic role of odor preprocessing by a shallow network of compartmentalized olfactory receptor neurons. Each compartment operates as a ratiometric unit for specific odor-mixtures. This computation arises from a simple mechanism: electrical coupling between two differently-sized neurons. We demonstrate that downstream synaptic connectivity is shaped to optimally leverage amplification of a hedonic value signal in the periphery. Furthermore, peripheral preprocessing is shown to markedly improve novel odor classification in a higher brain center. Together, our work highlights a far-reaching functional role of the sensory periphery for downstream processing. By elucidating the implementation of powerful computations by a shallow network, we provide insights into general principles of efficient sensory processing algorithms.

## Introduction

Animals navigating in natural environments process rapidly changing stimuli [1, 2], that are high-dimensional (i.e., consist of multiple components). Different organisms have adopted a broad range of strategies for detecting relevant stimuli within such sensory environments, to generate appropriate behavioral outputs. Major efforts in neuroscience are dedicated to understanding how the mammalian brain, utilizing elaborate and highly plastic neural architectures, solves these problems [3, 4]. The compact nervous systems of insects offer an opportunity to discover alternative strategies for efficient sensory processing. In particular, the conserved compartmentalization of primary sensory neurons [5–8] suggests that hard-wired peripheral preprocessing of sensory information enables organisms to effectively utilize shallow neural architectures for implementing powerful sensory processing algorithms.

In *Drosophila melanogaster*, the repertoire of olfactory receptor neurons (ORNs) in the periphery is genetically determined, and their compartmentalization into sensory hairs (‘sensilla’) is stereotyped [9,10]. ORNs housed in the same sensillum exert non-synaptic (‘ephaptic’) lateral inhibition [5]. We previously showed that these electrical interactions can affect odor-guided behaviors [5, 11, 12]. In natural environments, fruitflies encounter high-dimensional and transient odor stimuli [2], that are first processed by this conserved array of ORNs. The olfactory system of *Drosophila* provides a unique opportunity to understand the significance of neuronal compartmentalization for processing natural stimuli, because of the extensive information on its neuroanatomy [11, 13–17], and the rich theoretical work on how fly brains implement olfactory processing algorithms [18–20].

Olfactory signals from the periphery are transmitted to the Antennal Lobe (AL), from which inputs are routed along two main pathways (Figure 1A). Previous work has shown that connections from the AL to Lateral Horn (LH) mediate innate odor-guided behaviors [15, 21]. So far, behavioral experiments have mainly focused on responses to low-dimensional stimuli (typically, a single odorant, e.g., ethyl acetate, 2-hexanol [21], CO_2_ [22, 23]; or odor-guided behaviors mediated by a few glomeruli, e.g., apple cider vinegar [24]). However, it is unclear how behavioral responses to innately meaningful, high-dimensional odors are generated, and how the specific structure of AL-to-LH connections facilitates these computations. In contrast, AL to Mushroom Body (MB) connections are largely random [14, 25], and MB responses are sparse [26]. Architectures with random projections and sparsification are thought to facilitate robust learning of arbitrary stimulus-response associations, by increasing the linear separability of high-dimensional stimuli [18–20]. The hypothesis of enhanced linear separability in the MB relies crucially on the assumption of *clustered* neuronal responses in the AL [18], i.e., odors that elicit the same behavioral response correspond to similar AL population activity patterns. This assumption is inconsistent with empirical data showing, for example, that AL representations of different odors at the same concentration are often closer than that of the same odor at different concentrations [27, 28].

**Figure 1.**
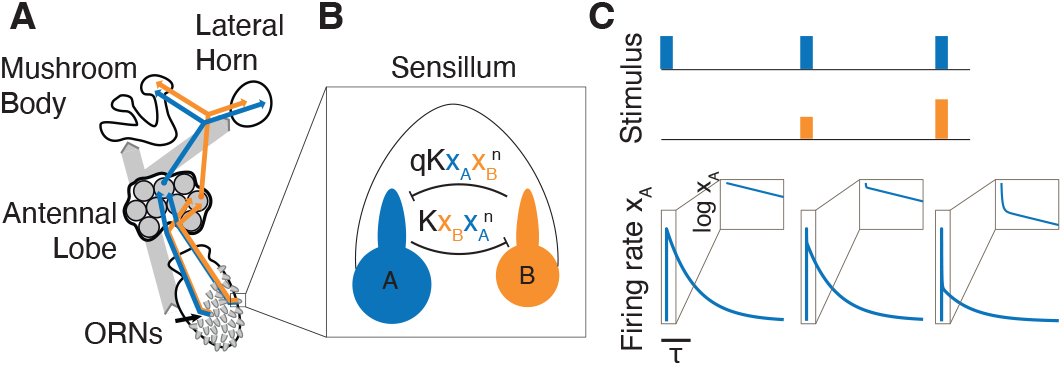
Nonlinear model of peripheral ephaptic interactions. (A) Illustration of olfactory information flow in fruitflies. (B) Peripheral signal preprocessing is mediated by ephaptic interactions between co-housed ORNs, wherein the neuronal firing-rates (*x*_*A*_, *x*_*B*_) are nonlinearly coupled. Model parameters *K, q, n* denote interaction strength, asymmetry and nonlinearity, respectively. (C) Analytical solutions of the response of neuron *A* (bottom) following offset of three different stimuli (top). Here, the strength of the *A* odorant (blue) is constant, while the strength of the *B* odorant (orange) increases. Activating neuron *B* leads to suppression of neuron *A* response. Insets: Firing-rate response on log scale illustrates a two-phase decay of the response to 0.

To address these fundamental questions of sensory processing in compact nervous systems, we focused on the role of lateral inhibition between compartmentalized ORNs [5,11] that transforms olfactory signals *before* they arrive at the AL [29]. We hypothesized that this peripheral preprocessing step enables efficient downstream integration of olfactory inputs.

## Results

### Lateral ephaptic inhibition between ORNs prepares olfactory signals for downstream processing

We constructed a simplified model of electrical (‘ephaptic’) inhibition between ORNs housed in the same sensillum (‘coupled ORNs’). The pairing of coupled ORNs is genetically determined [9], and their neuronal sizes exhibit stereotypical asymmetry [11,13]. The characteristics of the ephaptic inhibition each ORN exerts on its neighbor are influenced by the relative physical dimensions of the coupled ORNs [11, 30]. For tractability, here we focused on the typical case of a sensillum with two ORNs (Figure 1B). Each ORN’s label corresponds to its relative size (*A*, large; *B*, small). The time evolution of ORN *A*’s firing-rate (denoted *x*_*A*_) is modeled as,

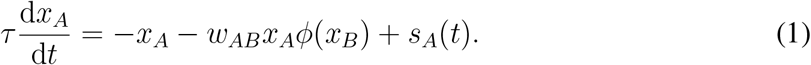

The three terms on the right respectively model a decay to baseline with timescale *τ*, nonlinear ephaptic inhibition by neuron *B*, and the time-dependent concentration of neuron *A*’s preferred odorant (i.e., the stimulus). The inhibition term contains two essential nonlinearities: 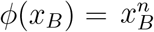 imposes a soft threshold for neuron *B*’s activation to influence its neighbor [31]; and the factor of *x*_*A*_ implies that the degree of inhibition experienced by neuron A depends on its own level of activation [5]. The evolution of *x*_*B*_ is given by replacing *A* with *B* in Eq. (1). We defined the asymmetry of interaction strengths as *q* = *w*_*BA*_*/w*_*AB*_. As neuron B is smaller, *q* ≤ 1, with smaller values of *q* implying more asymmetry in ephaptic interactions. We expect *q* to be a key sensillum-type specific parameter that tunes the asymmetry of ephaptic inhibition. Thus, multiple ‘copies’ of Eq. (1), with different values of *q*, represent an array of ORNs housed in different sensillum types that process high-dimensional olfactory stimuli.

Insects move through turbulent environments and encounter odorants sparsely in time [2, 32]. Therefore, we modeled stimuli as an instantaneous rise in concentration of the two odorants (*s*_*A,B*_(*t*) = *S*_*A,B*_*δ*(*t*)), which sets the ‘initial-condition’ of the firing rates, *x*_*A,B*_(*t* = 0) = *S*_*A,B*_ (Methods, [33]). We derived exact analytical solutions for the two-phase transient decay of *x*_*A,B*_(*t*) following stimulus ‘offset’ (Methods). The initial decay of *x*_*A*_ depends strongly on the concentration of odorant *B* due to ephaptic interactions (Figure 1C). After this initial phase, the firing rates decay exponentially, with a negligible effect of the coupling. Thus, our model’s complex transients contain information about the odor stimulus, consistent with experimental data [34].

Previous studies have demonstrated that ORN activity carries information about the behavioral valence of incoming stimuli. In particular, large ORNs (denoted ‘*A*’) are positive—their activation tends to trigger odor-guided behaviors such as attraction, egg-laying and courtship. Conversely, small ORNs (denoted ‘*B*’) are typically negative, antagonizing the behavioral output of the neighboring *A* neuron (ref. [12] and references therein). Based on these findings, we hypothesized that the computational role of ephaptic inhibition between behaviorally antagonistic ORNs is to enhance the net valence signal extracted from countervailing stimuli. The amplification of this valence signal could trigger an appropriate behavior by downstream regions.

Our model predicts that odor stimuli satisfying the linear scaling relationship *S*_*A*_ = *q*^1*/n*^*S*_*B*_ induce equal and opposite inhibition between the ORNs. Thus, we assumed that these stimuli are *neutral* (Methods). We defined valence amplification as the normalized net valence conveyed by the firing rates of the coupled ORNs,

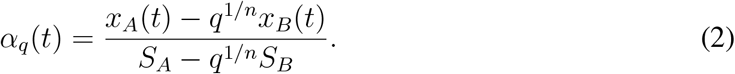

The solution to Eq. (1) shows that ephaptic interactions transiently amplify valence, i.e., *α*_*q*_(*t*) rises above 1 before decreasing to 0 at long times (Figure 2A,B). We note that the factor of *q*^1*/n*^ in Eq. (2) is necessary; any alternate (e.g., unweighted) definition of valence results in misidentified valence for a significant region of stimulus space (Figure S1, see Methods). This implies that to accurately detect stimulus valence based on the ORNs’ firing-rates, downstream readouts of ORN activity must ‘reweigh’ their inputs to calibrate the valence signal to neutral. More precisely, we predict that readout-weights are aligned to the angle tan^−1^(*q*^−1*/n*^), which depends on the interaction parameters for each sensillum. Ephaptic inhibition between coupled ORNs induces curvature of firing-rate trajectories as they decay to zero. Depending on the relative strengths of the odorants, the trajectories curve towards axes representing the activation of either the *A* or *B* neuron (Figure 2C). Ephaptic interactions therefore facilitate the discrimination of similar odor mixture ratios, based on the transient ORN responses. Both discriminability and valence amplification are maximal for odor stimuli with weak (i.e., close to neutral) valence (Figure 2B,C, dashed lines).

**Figure 2.**
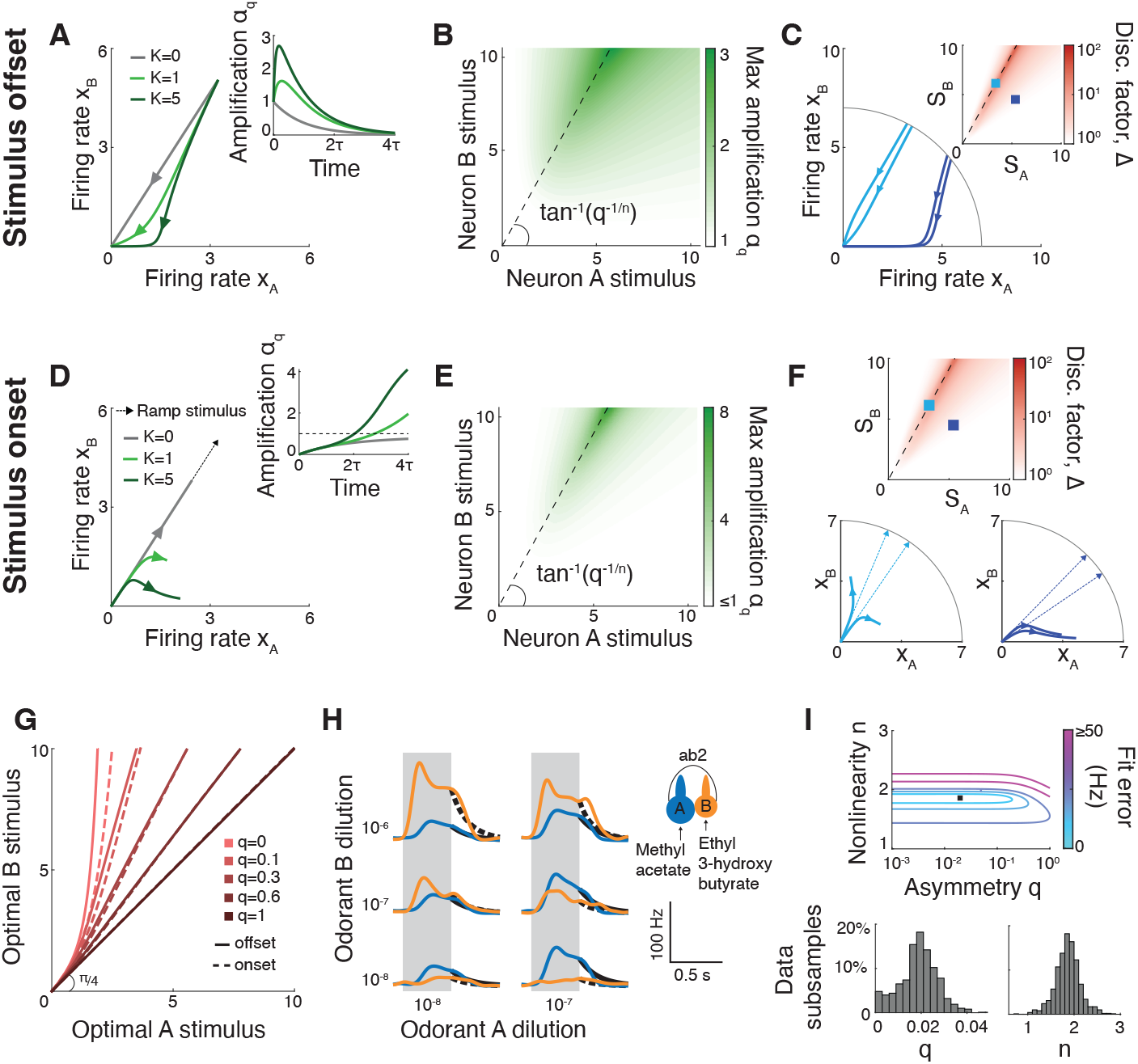
Ephaptic interactions transiently amplify odor valence. (A) Firing-rate trajectories following stimulus offset for different values of interaction strength *K*. Inset: Valence amplification *α*_*q*_(*t*) corresponding to the trajectories. In the presence of ephaptic interactions, valence is transiently amplified (*α*_*q*_(*t*) *>* 1). (B) Magnitude of valence amplification following stimulus offset [peak of *α*_*q*_(*t*)] as a function of the stimuli *S*_*A*_, *S*_*B*_. Amplification is stimulusspecific: maximal amplification is achieved for stimuli close to neutral (dashed line). (C) Example firing-rate trajectories for two sets of stimuli: weak valence (light blue) and strong valence (dark blue). Following stimulus offset, the light blue trajectories are separated in time, while the dark blue trajectories collapse onto each other and lose their initial separation. Inset: discrimination factor Δ (Methods) for different combinations of *S*_*A*_, *S*_*B*_. Large Δ indicates that the temporal evolution of the firing-rates can be used to discriminate between similar stimuli. Dashed line: neutral stimuli. (D) Same as (A), for stimulus onset (stimulus ramping up linearly in time, dashed black line). In the presence of ephaptic interactions, *α*_*q*_ attains a final value greater than 1, indicating valence amplification. Here, the duration of stimulus ramp is *T* = 4*τ* . (E) Same as (B) for stimulus onset. Valence amplification is maximal for stimuli close to neutral valence (dashed line). (F) Plot shows the discrimination factor Δ (top) and example firing-rate trajectories for two sets of ramp stimuli (bottom, dashed lines). Light blue trajectories corresponding to stimuli with weak valence as separated in time, whereas the dark blue trajectories (strong valence) lose their initial separation. This is summarized by the Δ values of the corresponding stimuli, which is maximal for stimuli close to neutral (dashed line). (G) Curves of optimal stimuli (with largest Δ, see Methods) for stimulus offset (solid lines) and stimulus onset (dashed lines) scenarios. The optimal stimuli depend strongly on the interaction asymmetry *q*. (H) Electrophysiological recordings of ORNs in the ab2 sensillum, probed with different odorant mixtures (data from ref. [11]) that were fit to the model (Methods). Gray shade: odor presentation duration. Black lines (solid: ab2A, dashed: ab2B) show model predictions following stimulus offset. (I) Fitting error for ab2 as a function of interaction asymmetry and nonlinearity parameters (top). Best fit values, *q* = 0.019 ± 0.007, *n* = 1.9 ± 0.26 (mean ± S.D., black square). S.D. obtained by subsampling trials (bottom). For (A-G), *q* = 0.3, *n* = 2, *K* = 1 unless noted otherwise.

As animals traverse through odor plumes, odorant concentrations follow complex temporal dynamics [2], which may not be captured by the pulse shape we have assumed so far (Figure 1C, Figure 2A-C). To show that our conclusions for stimulus offset apply in more general cases, we extended our analysis to the scenario of non-instantaneous stimulus onset. Indeed, in response to linear ramp stimuli of duration *T*, where *s*_*A,B*_(*t*) = *S*_*A,B*_*t/T*, transient ORN responses also exhibit amplified odor-valence (Figure 2D,E) and enhanced discriminability (Figure 2F).

Notably, our model suggests that the interaction asymmetry, specific to each sensillum type, determines which odorant ratio is ‘optimal’, i.e., has maximal amplification and discriminability. As *q* decreases, optimal stimuli are biased towards the negative valence odorant (Figure 2G, Methods). Moreover, optimal stimuli computed for stimulus offset and onset are in good agreement, for all values of the asymmetry *q* (Figure 2G, compare solid and dashed lines). This underscores the function of each sensillum type as a ratiometric processing unit. This unit of coupled ORNs could potentially extract behaviorally relevant olfactory information from odor mixtures with complex temporal dynamics of individual odorant concentrations.

We validated the model by fitting it to electrophysiology data from the ab2 sensillum (Figure 2H; assuming steady-state is reached within the 0.5 s stimulation, see Methods). The resulting asymmetry parameter *q* = 0.019 ± 0.007 ≪ 1 (Figure 2I, Figure S2) indicates a strong asymmetry in interactions between the ab2 ORNs, which is consistent with the large morphometric disparity measured for these neurons [13].

### Antennal lobe to Lateral Horn connectivity is structured to read out amplified odor-mixture valence

We hypothesized that the Lateral Horn (LH), the primary brain region mediating innate olfactory behaviors [21, 35], takes advantage of ephaptic interactions at the periphery to robustly compute the valence of high-dimensional odor stimuli. To do so, Lateral Horn neurons (LHNs) must asymmetrically weigh inputs from coupled ORNs to compute the amplified stimulus valence [numerator of Eq. (2)]. Given that *q <* 1, LH readout-weights for negative-valence ORNs are expected to be smaller. To test this prediction, we analyzed AL-to-LH projections in the Hemibrain connectome [16]. For simplicity, our analysis did not consider lateral interactions within the AL, or feedback connections (i.e., LH-to-AL, see Methods).

We focused on uni-glomerular Projection Neurons (uPNs), each of which constitutes a relay from a single ORN-type, thereby allowing us to directly examine ORN-specific signatures of projection asymmetry. There are one to three *types* of uPNs per glomerulus, which differ by their cell body locations and connectivity patterns [16, 36]. Our analysis showed that different types of uPNs from the same glomerulus had LH connectivity that was nearly as dissimilar as that of uPNs from different glomeruli (Figure S3). Therefore, we analyzed distinct uPN-types separately. LHN-types are also defined based on their location and other anatomical features [16]. As neurons of the same type have highly overlapping projection patterns (Figure S3), we considered a connection between each uPN-type and LHN-type to be a distinct information channel. This resulted in a connectivity matrix with 71 presynaptic uPN-types and 673 postsynaptic LHN-types (Methods). The weight of each connection was obtained by summing over the synapse counts between all individual neurons of the corresponding uPN- and LHN-types ([16], Methods). Henceforth, ‘uPN’ and ‘LHN’ refer to neuronal *types*, unless noted otherwise.

To identify salient connectivity features, it is important to compare the connectivity matrix to shuffles that preserve global properties of the connections [14, 19, 25]. We thus generated shuffled connectivity matrices that preserve the total number of incoming and outgoing connections per neuron in the original data (Figure S4). In addition to the connection frequencies, we preserved either incoming connection *weights* for each LHN (type-1 shuffles), or outgoing weights for each uPN (type-2 shuffles, see Methods and Figure S5). The two types of shuffles allow us to control for overalls biases in connection weights of LHNs (type-1) and uPNs (type-2). Principal Component Analysis (PCA) of the empirical and shuffled matrices revealed that the first few principal components (PCs) of the data explain a significantly larger amount of variance than expected by chance (Figure 3A, Figure S6, in contrast to ref. [14] where PCA of AL-to-MB connections revealed no such PCs). Thus, these PCs represent structured connections from uPNs to LHNs, with the leading PC (PC1) describing the most prominent pattern of AL-to-LH readouts. The PC1 coefficient of each uPN quantifies its weight in the aforementioned readout pattern.

**Figure 3.**
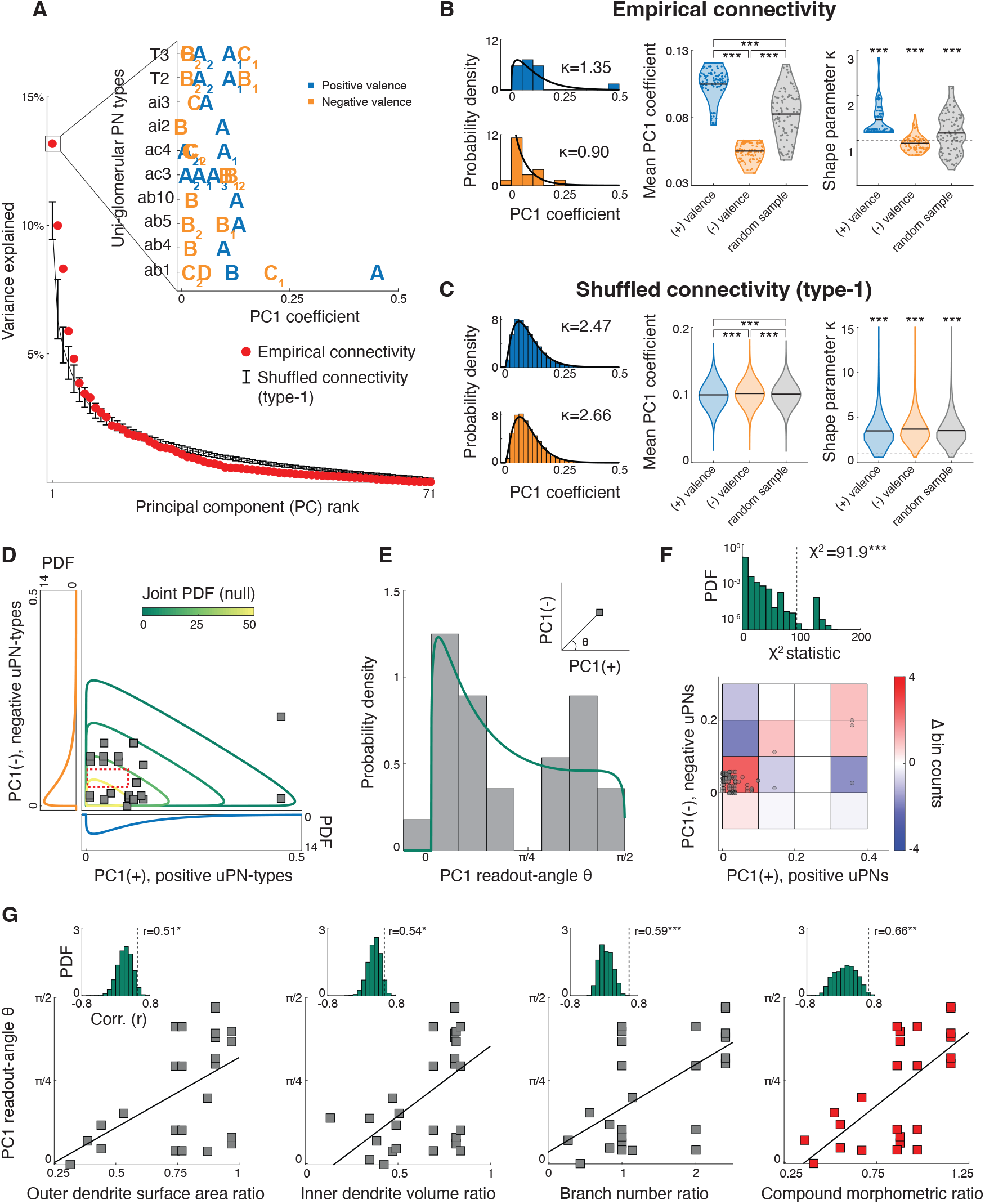
Antennal Lobe (AL) to Lateral Horn (LH) projections are structured to compute amplified stimulus valence. (A) Variance explained by Principal Components of the empirical AL-to-LH connectivity matrix (red) and type-1 shuffles that preserve incoming connection weights for each LHN-type (black, error bars indicate 95% confidence intervals). Inset: PC1 coefficients of uPN-types corresponding to behaviorally antagonistic coupled ORNs [12], grouped by sensilla. Subscripts label multiple uPN-types per ORN (Table S1). (B) Left: Distributions of PC1 coefficients for positive(*n* = 14) and negative-valence (*n* = 16) uPN-types, and the best fit Gamma distribution (black) for the empirical connectivity. See Methods for details of fitting procedure, including handling of one negative PC1 coefficient. Right: The average PC1 coefficient and Gamma distribution shape parameter (*κ*) obtained from 100 subsamples of positive- and negative-valence uPN-types (blue and orange, respectively), as well as 14 randomly selected uPN-types (gray, 100 samples). Data points indicate individual values obtained from each subsample, and black lines indicate the mean. Positive- and negative-valence uPN-types have average PC1 coefficients that are respectively greater and smaller than that obtained for random samples. Further, the average PC1 coefficient for positive uPN-types is significantly larger than that for negative uPN-types. The distribution of PC1 coefficients for negative uPN-types corresponds to a shape parameter *κ <* 1, indicating a mode at zero. The PC1 coefficient distributions for positive uPN-types and random samples correspond to *κ >* 1. ^**^*P <* 0.01, ^***^*P <* 0.001, one-sided Student’s t-test. (C) Same as (B) for PC1 coefficients obtained from type-1 shuffled connectivity matrices. Here, the mean PC1 coefficient for positive uPN-types is slightly smaller than that of negative uPN-types and random samples. This difference, though statistically significant, is two orders of magnitude smaller than that seen in the empirical data. Here, the PC1 coefficient distributions for both positive and negative uPN-types, as well as random samples correspond to shape parameters *κ >* 1. These results demonstrate a lack of bias between positive and negative uPN-types in the shuffled connectivity. Plots were generated using 10^4^ shuffles, with 100 subsamples each (Methods). (D) PC1 coefficients of pairs of uPN-types postsynaptic to coupled, behaviorally antagonistic ORNs (squares, *n* = 25). Also shown are contour lines of the null distribution, obtained by assuming PC1 coefficients for the pairs are statistically independent (Methods). Based on the null distribution, we expected to find data points within the red rectangle. The absence of data points suggests a poor match between the null and empirical distributions. We note that the marginal PC1 coefficient distributions in (B,D) differ. In (B), each uPN-type is counted once, while the marginal distributions in (D) are constructed by counting uPN-types once for each pair they participate in. (E) Inset: Calculation of PC1 readout-angle (*θ*) for a uPN-type pair. Empirical (gray) and null (green) distributions of *θ* for the PC1 coefficient pairs in (D) (see Methods). PC1 readout-angles close to *π/*4 (symmetric readouts) are noticeably absent in the data, which is poorly explained by the null distribution. (F) Results of Pearson’s chi-squared test of independence for PC1 coefficients of *individual* uPNs. Gray circles: PC1 coefficients of pairs of uPNs postsynaptic to coupled, behaviorally antagonistic ORNS (*n* = 124). Colors indicate the difference between empirical and expected number of data points in each bin, which is used to calculate the *χ*^2^ statistic (top; see Methods, Figure S8 for other bin-sizes). The *χ*^2^ statistic for the data is significantly larger than that obtained for pairs of PC1 coefficients generated under the assumption of statistical independence (green histogram), indicating significant statistical dependencies in the empirical distribution (G) PC1 readout-angles of paired uPN-types versus ratios of three dendritic measurements of the corresponding ORN pairs, and a compound morphometric ratio (Methods). The quantities are positively correlated. Insets: Analogous correlation values for random pairs of positiveand negative-valence ORNs are significantly smaller than that for empirical ORN pairs (Methods). ^*^*P <* 0.05,^**^ *P <* 0.01,^***^ *P <* 0.001, *P* -values computed using distribution percentiles.

Based on the available valence information for ORNs [12], we identified 30 uPNs that were postsynaptic to coupled, behaviorally antagonistic ORNs (*n* = 14 positive valence, *n* = 16 negative valence; Figure 3A, inset; see Table S1). The mean PC1 coefficient for negative-valence uPNs is significantly smaller than that of positive-valence uPNs (Figure 3B, positive-valence uPNs: 0.105 ± 0.012, negative-valence uPNs: 0.055 ± 0.006). We found that these values were significantly different than the mean PC1 coefficient of a randomly chosen sample of 14 uPNs (0.083 ± 0.018, Figure 3B), indicating that PC1 coefficients of positiveand negativevalence uPNs are distributed differently. Next, we fitted a Gamma distribution to each set of PC1 coefficients separately (Figure 3B). The distribution mode for negative-valence uPNs is 0, while for positive uPN and random samples it is strictly positive. This is indicated by the shape parameters of the corresponding Gamma distributions (Figure 3B) – positive uPNs: 1.45 ± 0.38, negative uPNs: 0.93 ± 0.18, random sample: 1.16 ± 0.44. Taken together, negative-valence uPNs are biased towards smaller PC1 coefficients. In contrast, this bias is negligible for shuffled connectivity matrices (Figure 3C, Figure S6). This implies that the predicted asymmetry in readout-weights indeed arises from structured AL-to-LH connectivity, and cannot be explained simply by biases in the frequency of connections or connection weights.

Our model further predicts that in order to compute the amplified valence signal, the asymmetry in LH readout-weights of coupled ORNs must depend on the interaction asymmetry (*q*). We identified 25 uPN pairs in the connectome that correspond to behaviorally antagonistic ORNs from the same sensillum [12]. Indeed, the connectivity of 16/25 pairs (64%) differed significantly between the data and shuffles (Figure S7), indicating dependencies between LHN targets of these uPN pairs. If AL-to-LH connectivity indeed reflects ORN coupling at the periphery, the readout-weights of pairs of behaviorally antagonistic, coupled ORNs would be distributed differently than that of random pairs of positive- and negative-valence ORNs. We thus compared the joint distribution of the 25 pairs of PC1 coefficients to the null distribution, obtained by independently sampling from the marginals of positive- and negative-valence uPNs (Methods). Notably, there is a region containing no data points (Figure 3D, red box), suggesting that the empirical data is not fully explained by the null distribution. We further computed readout-angles of the uPN pairs, which exhibit a marked discrepancy between the empirical and the null distributions (Figure 3E). Angles close to *π/*4, corresponding to symmetric readouts, are noticeably not represented in the distribution. However, we were unable to rule-out the null hypothesis in Figure 3D,E in statistical tests, likely due to the relatively small number of uPN-types corresponding to ORNs with known behavioral valence [12].

Our model predicts *global biases* in readout-weights that apply at the level of input channels, i.e., uPN *types* post-synaptic to coupled ORNs. However, the predicted *dependencies* between readout-weights of coupled ORNs can be examined at the level of individual neurons. Therefore, we analyzed PC1 coefficients of *individual* uPNs to test the null hypothesis with increased statistical power. Indeed, we found a statistically significant dependence between the empirical PC1 coefficient pairs (Figure 3F), and show that the distribution of PC1 readout-angles is significantly different from the corresponding null distribution (Figure S8). Together these statistical tests demonstrate that the readout weights of ORNs reflect their coupling in the periphery.

Next, we tested the hypothesis that AL-to-LH connectivity reflects asymmetrical coupling between ORNs by studying the relationship between the readout-angles and morphometric asymmetry of coupled ORN pairs [13]. If our hypothesis is correct, a positive correlation is expected between these quantities, since asymmetry in ephaptic interactions arises from ORN size disparities [Eq. (2), Refs. [11, 13, 30]]. We computed ratios of five dendritic measurements and two somatic measurements (Figure S9, ref. [13]) for each pair of coupled ORNs. Three of the five dendritic ratios, but none of the somatic ratios, had a significant positive correlation with PC1 readout-angles, when compared to the control: analogous correlation values obtained for random pairs of positive- and negative-valence ORNs (Figure 3G). To quantify the overall morphometric asymmetry between coupled ORNs, we further defined a *compound morphometric ratio* as a linear combination of these three ratios. The coefficients of the compound ratio were optimized to maximize its correlation with PC1 readout-angles (Figure 3G, Methods). The resulting correlation of *r* = 0.66 was highly significant relative to the control. Notably, the fact that somatic ratios were not positively correlated with the readout-angles (Figure S9) is consistent with the notion that ephaptic coupling arises from interactions between dendritic compartments [5, 11]. We did not find similar correlations between morphometry and readout-angles for secondary PCs of the AL-to-LH connectivity matrix (see Methods and Figure S10), suggesting that the aligned valence readout is achieved primarily by the most prominent projection pattern.

We note that the randomization procedure used as a control in Figure 3G generates random ORN pairs that have opposite valence, but do not correspond to the same sensillum. Therefore, the significance of the correlations between peripheral morphometric ratios and downstream readouts demonstrates that AL-to-LH projections reflect the compartmentalized structure of the olfactory periphery, *beyond* valence opponency. Based on our computational model of ephaptic interactions between coupled ORNs, we argue that the aligned readouts of the entire sensilla array allow for effective propagation of the transiently amplified valence signal extracted from *any* odorant mixture.

### Peripheral preprocessing enables odor classification in the mushroom body

Finally, we studied the functional impact of peripheral ephaptic inhibition on odor processing in the Mushroom Body (MB). AL-to-MB projections are largely random, and expansive: projection neurons (PNs) corresponding to ∼50 glomeruli relay information to 2000-2500 Kenyon Cells (KCs) in the MB [14, 25, 37]. KC responses are sparser than those of PNs [26]. Expansive and sparse projections—which increase linear separability of high-dimensional stimuli—was proposed as the key mechanism supporting novel odor classification in the MB [18, 20]. However, a crucial assumption in these studies is an initial clustering of stimuli in the input layer. This clustering implies that perceptually similar odors elicit similar AL glomerular responses (Figure 4A), which is not consistent with empirical data obtained from fruitflies and other insects [27, 28].

**Figure 4.**
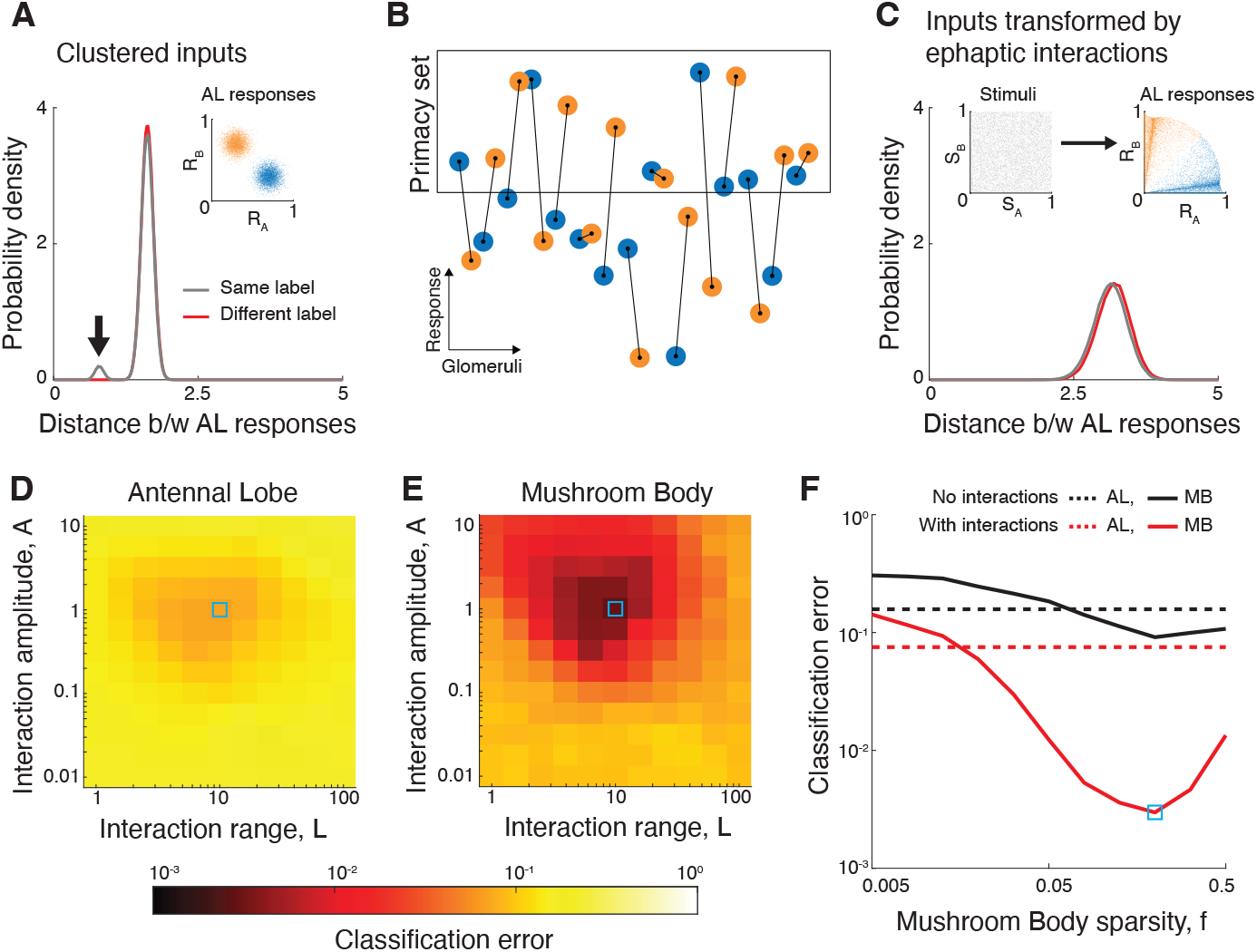
Ephaptic interactions improve odor classification in the Mushroom Body. (A) Distance between stimuli (in *N* = 50 dimensional response space) with the same (gray) or different (red) labels under the assumption of clustered AL responses. Stimuli with the same label are closer, as indicated by the peak in the gray distribution (black arrow). Inset: 2D illustration of clustered AL responses. (B) Model of primacy coding in the Antennal Lobe. Glomeruli with the largest responses constitute the primacy-set for the stimulus, and determine the stimulus label. Black lines indicate glomeruli correspondingg to ephaptically coupled ORNs. (C) Same as (A) for uniformly distributed stimuli that are transformed by ephaptic interactions and assigned a label based on the primacy set. Stimuli with the same or different labels have similar distance distributions (See also Figure S11). (D) Classification error of a Support-Vector Machine (SVM) based on AL responses (Methods), as a function of the ephaptic interaction parameters. The optimal parameter combination leads to modest improvements. (E) Same as (D) for MB responses (i.e., following random expansion and sparsification, sparsity *f* = 0.2). Here ephaptic interactions result in marked improvements (note the logarithmic color-scale). SVM classification error as a function of the sparsity of MB responses *f* . In the absence of ephaptic interactions (black curves), expansion and sparsification in MB does not result in a significant improvement in classification performance. See Methods for simulation details.

Given that ephaptic interactions can transiently separate ORN responses based on valence (Figure 2C,F), we hypothesized that preprocessing of AL responses by peripheral interactions may facilitate odor classification in the MB—without requiring explicit clustering. To test this hypothesis, we simulated the response of *N* = 50 ORNs selective to *N* uniformly distributed private odorants, each having positive- or negative-valence. There were *N/*2 pairs of ORNs with opposite valence that were ephaptically coupled in the simulations (Figure 4B and ref. [5, 12]). For simplicity, we analyzed ORN responses at a particular ‘snapshot’ in time. Ephaptic interactions were modeled as shifts in ORN responses relative to their initial values at *t* = 0 set by an odorant concentration ‘pulse’. To calculate the response shift, the interactions were described by two parameters: interaction amplitude *A* and interaction range *L* (Methods). The range refers to the distance (in stimulus-space) from neutral stimuli for which inhibition is significant. The direction and magnitude of the shifts depend on the net stimulus valence, similar to our time-dependent model (Eq. (1), Figure 2A-F, Figure S11). Glomeruli were assumed to be one-to-one relays of ORN activity, as in our connectomic analysis.

We studied the classification of odor stimuli into two categories (e.g., appetitive or aversive), as in [18]. In contrast to previous studies [18], we did not assume that glomerular responses for each category are clustered. Instead, stimuli were assigned category labels based on the principle of *primacy coding* in the AL (Figure 4B). Primacy coding implies that the identity of an odor is encoded by the glomeruli that respond most strongly to it (i.e., its ‘primacy-set’ which is concentration invariant [38, 39]). Thus, we labeled a stimulus as +1 if the number of positive-valence glomeruli was greater than negative-valence glomeruli in its primacy-set, and −1 otherwise (Figure 4C, inset). Our analysis showed that ephaptic interactions impose a subtle structure on the distribution of glomerular responses, compared to the case of clustered inputs (Figure 4A,C, Figure S11).

We compared classification performance in the AL using responses computed with or without ephaptic inhibition, and identified a region of *A, L* values where interactions lead to modest improvements (Figure 4D). Remarkably, we found that random expansion of the preprocessed AL responses to the MB layer, followed by sparsification of the responses ([18], Methods) resulted in a significant improvement in classification performance (Figure 4E,F). In contrast, random expansion of AL responses calculated without ephaptic interactions resulted in no such improvements (Figure 4F). Thus, ephaptic interactions can render classification following random expansion and sparsification effective, despite the absence of a clustered input-layer. We further studied the case where glomeruli were partitioned randomly, rather than by valence. Stimulus labels were assigned based on which of the two partitions had a majority in the primacy-set. We found that ephaptic interactions moderately enhanced classification in this scenario as well (Figure S12), implying that information preprocessing by coupled ORNs can potentially facilitate learning of *arbitrary* stimulus–label associations in the MB.

## Discussion

Here, using a combination of mathematical modeling and analysis of experimental data, we have shown how peripheral interactions between ORNs work in coordination with the wiring diagram in the central brain to optimize complex sensory computations. Previously, anatomical wiring in fruitflies was shown to implement *specific* sensorimotor computations, for example, detection of two-dimensional visual motion within a restricted field of view [40], detection of coincident noxious stimuli [41], and evaluation of heading direction [42]. Here, using our model of *peripheral* preprocessing, we made predictions for *central* connectivity necessary to perform robust, valence-based classification of complex odors. By validating these predictions, we demonstrated how the wiring-diagram implements a *general purpose* computation: extracting the valence of arbitrary odors. Our study further suggests that a sparse and random expansion of sensory information [14,18,19] can enable classification, if AL responses are transformed by peripheral interactions. We have demonstrated that this scheme can be effective without clustering, i.e., without stimuli of the same category being contained within a radius [43]. Therefore, our study can inspire future research of engineered systems, to investigate other computationally inexpensive transformations of the input layer that can similarly enable classification by the random expansion and sparsification architecture [20, 44]. Such architectures could support powerful sensory computations without relying on costly training or metabolically inefficient hierarchical processing.

Our findings highlight the importance of peripheral processing for central sensory computations, which may have implications in biological systems beyond olfaction. Indeed, functionally similar lateral inhibition was reported between the R7 and R8 photoreceptors [7], which have different spectral sensitivity profiles in the *Drosophila* compound eye. Future work may reveal how such color-opponent preprocessing contributes to initiation of appropriate visually guided behaviors in flies.

## Acknowledgments

The authors thank J.W. Wang and D. Kleinfeld for useful discussions and comments on the manuscript. This work was supported by DARPA grant D21AP10162-00 (J.A.), DOE grant DE-SC0022042 (J.A.), NIH grants R21AI169343, R21DC020536, R01DC016466 (C.-Y.S.),an Innovative Research Grant from the Kavli Institute for Brain and Mind (UC San Diego) (awarded jointly to C.-Y.S., J.A.). P.P. thanks the UC San Diego Friends of the International Center, and UC San Diego GPSA for support.

## Author contributions

Developed model, P.P., J.A. with inputs from S.-T.W., C.-Y.S. Solved, analyzed and simulated model, P.P., J.A. Designed and performed data analysis, P.P. with inputs from J.A., S.T.W., C.-Y.S. Wrote paper, P.P., J.A. with inputs from S.-T.W., C.-Y.S. Acquired funding and supervised project, J.A., C.-Y.S.

## Methods, Supplementary Figures and Supplementary Table

### 1 Methods

#### 1.1 Reduced nonlinear model of a sensillum

We constructed the following nonlinear model of the responses of co-housed ORNs,

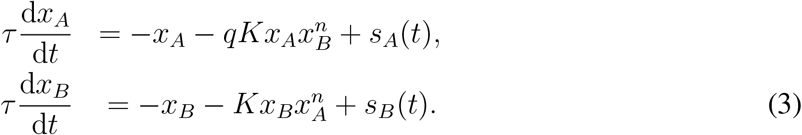

The parameters are introduced in the main text. To study the dynamical behavior of this model, we considered a pulse stimulus, *s*_*A,B*_(*t*) = *S*_*A,B*_*δ*(*t*).

##### 1.1.1 Analytical solution for transient decay

For *t >* 0 (after stimulus removal), Eq. (3) can be rewritten as,

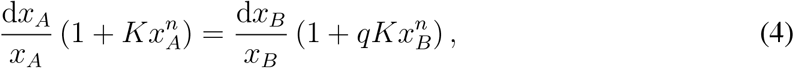

with the initial conditions *x*_*A,B*_(*t*) = *S*_*A,B*_. Since the leftand right-hand sides of Eq. (4) depend separately on *x*_*A*_ and *x*_*B*_, we integrated both sides,

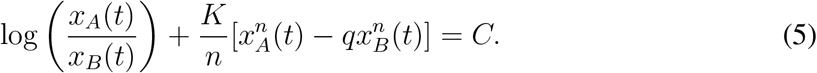

Importantly, *C* is a constant. Thus we have identified an invariant of the dynamics, determined by the stimulus,

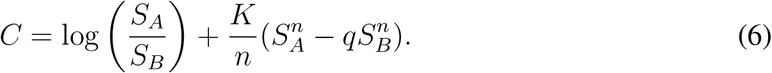

Next we defined *u*(*t*) = log[*x*_*A*_(*t*)*/x*_*B*_(*t*)], computed its derivative directly and substituted in *C*,

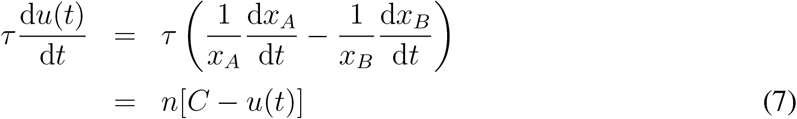

The solution of Eq. (7) is,

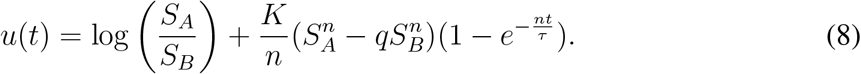

We then substituted the dynamical equation of *x*_*B*_ into Eq. (7) and rearranged the terms to get,

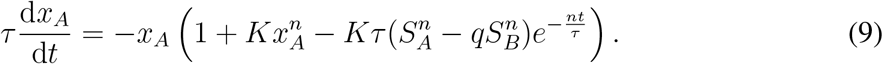

Eq. (9) involves only *x*_*A*_. One can check that its solution, and the solution to the analogous equation for *x*_*B*_ are,

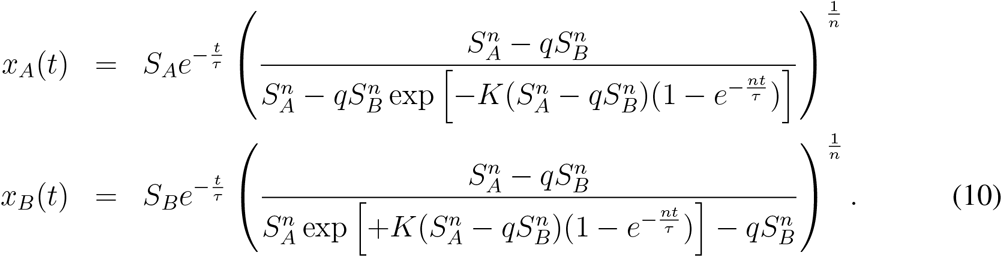

##### 1.1.2 Definition and solutions of valence amplification *α*_*q*_(*t*)

The solutions to the dynamical model Eq. (10) show that for stimulus mixtures satisfying 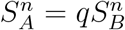, the firing-rates *x*_*A*_, *x*_*B*_ decay to zero along a straight line. In other words, for these stimuli, ORNs *A* and *B* receive equal inhibition from each other. Since large- and small-spike ORNs were shown to carry opposite valence signals [12], we interpreted stimuli satisfying this condition as *neutral*, and the weighted difference *S*_*A*_ −*q*^1*/n*^*S*_*B*_ as the net-valence of the stimulus mixture. We compared the net valence of the stimulus to that conveyed by ORN responses, by defining the transient valence amplification,

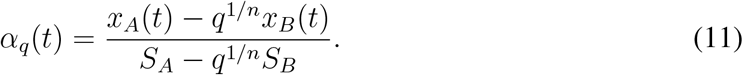

The formulae in Eq. (10) were used to evaluate *α*_*q*_(*t*) directly.

##### 1.1.3 Definition of the discriminability factor Δ

We calculated the time evolution of the angle of the firing rate trajectories *θ*(*t*) using Eq. (10),

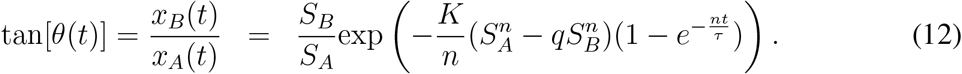

The initial value of *θ* is given by the relative strengths of the odorants, and at long times (*t* → ∞) it saturates to a value that depends on the stimuli as well as ephaptic interaction parameters. To understand if ephaptic interactions can aid in discrimination between responses to similar odor mixtures (i.e., stimuli with similar ratios *S*_*B*_*/S*_*A*_), we first defined stimulus sensitivity *σ*(*t*) as the partial derivative

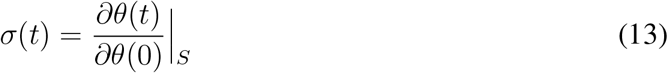

evaluated at fixed stimulus strength 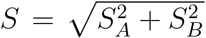. The partial derivative can be evaluated directly,

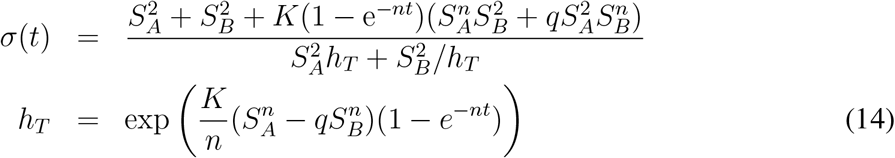

We further defined the discriminability factor Δ = *s*(*t*^*^), where *t*^*^ = argmax{*σ*(*t*)}. Thus, Δ ≥ *σ*(*t* = 0) = 1. Large Δ implies that two similar odor mixtures (of the same strength) can be easily discriminated by ORN responses. One sees from the denominator of *σ*(*t*) and the form of *h*_*T*_ that discriminability is maximal for neutral stimuli 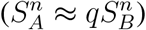, for which transient amplification is also maximal.

##### 1.1.4 Numerical solutions for stimulus onset

We considered the response of the model to a linear ramp stimulus, *s*_*A,B*_(*t*) = *S*_*A,B*_*t/T*, which attains its final value at *t* = *T* . In the absence of ephaptic coupling (*K* = 0), the analytical solution for the firing-rates is:

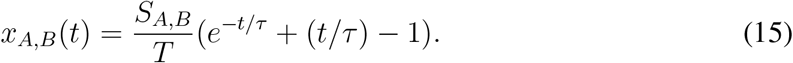

One sees that for fast ramping relative to the membrane time-constant (*T* ≪ *τ*), the firing-rates are very small *x*_*A,B*_(*T*) ≪ *S*_*A,B*_ at the end of the onset transient. For slow ramping (*T* ≫ *τ*), the firing-rates track the stimulus, *x*_*A,B*_(*T*) ≈ *S*_*A,B*_. To study the effect of ephaptic interactions on the response of the system, we focused on the intermediate regime, *T* ≈ *τ* . We numerically solved Eq. (3) for ramp stimuli to find the firing rate trajectories *x*_*A,B*_(*t*) in the presence of ephaptic inhibition (Figure 2D). Analogous to the case of stimulus offset, we defined a transient valence amplification,

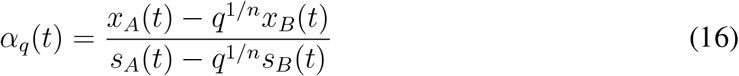

Transient amplification is initially 0 (*α*_*q*_(0) = 0), and it reaches a finite value at *t* = *T* when the firing-rates increase in response to the stimuli. In the presence of ephaptic interactions, the final value *α*_*q*_(*T*) *>* 1, indicating valence amplification (Figure S2A, inset). Valence amplification by ephaptic interactions is stimulus-specific (Figure 2E), and maximal for stimuli close to neutral. Ephaptic interactions enhance the discriminability of weakly positive and weakly negative odors (Figure 2F). Here, we computed the discriminability factor using our numerical solutions, 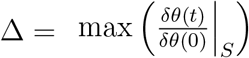 with *δθ*(0) = 0.01. Similar to the scenario of stimulus offset, there are optimal (i.e., maximally separated) stimuli for different values of asymmetry *q* (Figure 2G). Optimal stimuli for the case of onset responses are in excellent agreement with those obtained from analytical solutions for offset responses.

##### 1.1.5 Fitting model parameters to ab2 electrophysiological recordings

The ab2 sensillum consists of two olfactory receptor neurons, ab2A (Or59b) and ab2B (Or85a), that respond selectively to odorants Methyl acetate and Ethyl 3-hydroxy butyrate respectively. *In-vivo* single sensillum recordings were performed in response to a 0.5 s pulse of odorant mixtures at different dilution ratios [Figure 2H, 6 dilution ratios (‘conditions’), *n* = 9 trials each]. Data are taken from Ref. [11]. We assumed that the system is approximately at steadystate during odor presentation, with constant stimuli (*s*_*A,B*_(*t*) = *S*_*A,B*_). Substituting d*x*_*A,B*_*/*d*t* = 0 in Eq. (3), the nonlinear model reduces to,

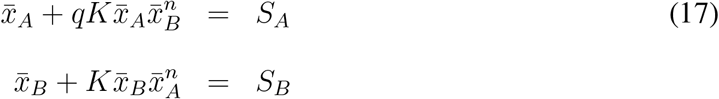

where 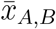 denotes the steady-state firing rates of the neurons, and *S*_*A,B*_ is the amplitude of the odor pulses (in units of Hz). In the absence of ephaptic interactions (*K* = 0), *S*_*A,B*_ represent the steady-state firing rates of the neurons in response to the odorants. We estimated the value of *S*_*A,B*_ for different odorant dilutions using ORN responses to single odorants (Figure S2A-C). We ignored initial transients, and calculated the steady-state responses 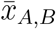 by averaging the firing rate from the time it reaches its peak until the end of odor presentation. Let 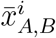 and 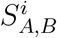 denote the steady-state responses and odorant amplitudes for condition *i*. We quantified the fitting error for different interaction parameters by,

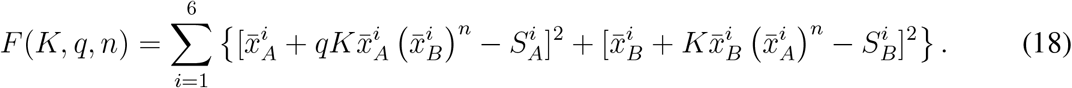

We utilized gradient descent methods to locate the minimum of *F* (*K, q, n*), which represents ephaptic interaction parameters that best fit ORN responses to all six odor mixtures (Figure S2E-G). A distribution of fitting parameters was generated by repeatedly dropping data from six random trials (on average, one trial from each condition; repeated 1000 times) in the calculation of 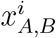 and then performing the fit (Figure 2I, Figure S2D).

#### 1.2 Analysis of AL-to-LH connections

##### 1.2.1 Analysis at the level of neuronal types

The Hemibrain connectome (Ref. [16], dataset version v1.2.1) contains 168 uniglomerular projection neurons (uPNs) and 1497 Lateral Horn neurons (LHNs). Based on similarities in morphology and connectivity features, uPNs and LHNs are grouped into 90 and 689 neuronal types, respectively. Out of these, all 19 uPN-types from the left hemisphere, and 16 LHN-types (14 right hemisphere, 2 left hemisphere) do not participate in any AL-to-LH connections. These neuron-types were removed from the subsequent analyses. Next, we quantified the difference between distinct types of uPNs that originate in the same glomerulus. Using the connectome data, we calculated the number of target LHN-types that any two uPNs have in common. We considered separately all uPN-pairs of the same type originating from the same glomerulus, uPN-pairs of different types but from the same glomerulus, and uPN-pairs corresponding to different glomeruli. Our analysis showed that on average, uPNs of the same type from the same glomerulus shared 75 postsynaptic LHN-types, while uPNs of different types from the same glomerulus shared only 39 postsynaptic LHN-types (Figure S3A). On the other hand, uPNs originating from different glomeruli had, on average, 32 LHN targets in common. The fraction of shared targets was 79%, 32% and 47% respectively (Figure S3B-D). This implies that LH connectivity of distinct uPN-types from the same glomerulus is nearly as dissimilar as that of uPNs from different glomeruli, motivating us to analyze the dataset at the level of uPN-types rather than glomeruli. We thus obtained a connectivity matrix with 71 uPN-types and 673 LHN-types. To calculate the weight of each connection between a particular uPN-type and LHN-type, we summed the synapse counts identified between the individual uPNs and LHNs of the respective types. Henceforth, ‘uPNs’ and ‘LHNs’ refer to neuronal *types* instead of individual neurons, unless noted otherwise.

##### 1.2.2 Connectivity shuffles with conserved uPNs and LHNs statistics

We constructed shuffled versions of the 71-by-673 dimensional empirical connectivity matrix by randomizing connections between uPNs and LHNs. We constrained the shuffles to ensure that each uPN and LHN participates in the same number of connections as in the original data, as in [14]. This preserves the frequency of incoming and outgoing connections in the empirical data (Figure S4), while removing any non-random, structured patterns of AL-to-LH connectivity. We implemented two types of shuffles, which differed in the constraint on connection weights that was imposed: **type-1** shuffles preserved the incoming connection weights for each LHN, while **type-2** shuffles preserved outgoing connection weights for each uPN (Figure S5). The only matrix that satisfies both constraints on connection weights is the empirical connectivity matrix itself. The two types of shuffles allow us to probe the significance of different properties of the empirical connectivity matrix. The variance of outgoing connection weights for type-1 shuffles is smaller than that of the empirical distributions (compare Figure S6A,B). This means that uPNs with substantially smaller or larger outgoing connection weights relative to the mean are not represented in type-1 shuffles, which allowed us to probe the consequences of biases in uPNs. Indeed, we found that the leading PCs in the empirical data explain more variance than expected from type-1 shuffles (Figure 3A). On the other hand, compared to the empirical data, the incoming connection weight distribution for type-2 shuffles is not heavy-tailed (i.e., ‘outlier’ LHNs receiving strong inputs are not represented, compare Figure S5D,F). Therefore, by comparing the empirical connectivity matrix with type-2 shuffles, we can isolate the consequences of perturbing biases in incoming weights for each LHN. Previous work [19] has shown that analogous statistics of incoming AL-to-MB connections are important for information processing. Again we found that leading PCs explain substantially more variance in the data compared to type-2 shuffles (Figure S6).

##### 1.2.3 Fitting Gamma distribution to PC1 coefficients

We analyzed the distribution of PC1 coefficients of uPNs carrying positiveand negativevalence signals (Table S1, Figure 3B). We fitted a Gamma distribution to each set of PC1 coefficients using maximum-likelihood estimation. A Gamma distribution has two parameters: scale (*θ*) and shape (*κ*), and its probability density function is given by

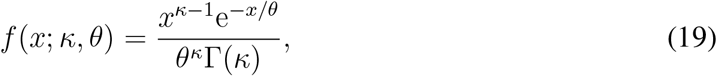

for *x* ≥ 0. Here, Γ is the Gamma function and distribution mean is given by *κθ*. In the empirical data, one uPN has a small, negative PC1 coefficient (-0.0006, small relative to the distribution mean 0.057). We replaced that value with 0 to obtain a valid fit for the Gamma distribution (Figure 3B, distribution of PC1 coefficients for negative valence uPNs). An alternate procedure, wherein we first shifted all PC1 coefficients by the negative coefficient (such that the smallest coefficient was now 0), and then fit the gamma distribution, yielded a similar result (not shown). All the subsequent conclusions did not depend on the choice of fitting procedure.

We subsampled negative- and positive-valence uPNs by dropping 10% of PC1 coefficients (Figure 3B, repeated 100 times). As a control, we also generated 100 samples of 14 randomly chosen uPNs. For each subsample and random sample, we calculated the mean PC1 coefficient and fit a Gamma distribution. We found that the mean PC1 coefficient of negative uPNs is smaller than that of positive uPNs (*P <* 0.001 from one-sided Student’s t-test). Positive- and negative-valence uPNs have mean PC1 coefficients that are respectively greater and smaller than that obtained for random samples (one-sided Student’s t-test, *P <* 0.001). Additionally, the shape parameter of the Gamma distribution *κ <* 1 for negative uPNs (one-sided Student’s t-test, *P <* 0.01), while *κ >* 1 for positive uPNs and random samples (one-sided Student’s t-test, *P <* 0.001). The shape parameter controls the distribution mode: for *κ <* 1, the mode is equal to zero, while for *κ >* 1 the mode is strictly greater than zero.

Next, we repeated the above procedure for shuffled datasets as a control (Figure 3C, Figure S6). In contrast to the empirical data, here, the mean PC1 coefficient for negative uPNs was slightly *larger* than positive uPNs and random samples– for type-1 shuffles: ⟨PC1⟩_+_ = 0.100 ± 0.017, ⟨PC1⟩_−_ = 0.102 ± 0.016, ⟨PC1⟩_rand_ = 0.101 ± 0.016; for type-2 shuffles: ⟨PC1⟩_+_ = 0.005 ± 0.014, ⟨PC1⟩_−_ = 0.005 ± 0.006, ⟨PC1⟩_rand_ = 0.005 ± 0.010. The differences in the mean PC1 coefficients for the shuffles, though statistically significant, are roughly two orders of magnitude smaller than that seen for the empirical data. Notably, type-2 shuffles have a significant fraction of negative PC1 coefficients. Since Gamma distributions have a positive support, we could not estimate shape parameters for these coefficients. For type-1 shuffles, the shape parameter *κ >* 1 for both positive and negative uPNs, and random samples (one-sided Student’s t-tests, *P <* 0.001, *κ*_+_ = 3.53 ± 1.76, *κ*_−_ = 3.72 ± 1.59, *κ*_rand_ = 3.55 ± 1.54). This indicates a mode strictly greater than zero for all three distributions. Therefore, we conclude that the PC1 coefficient distributions of positive and negative valence uPNs are similar for shuffled datasets, lacking the systematic bias observed in the empirical data.

##### 1.2.4 Null distribution for PC1 coefficient pairs

We calculated the joint distribution of PC1 coefficients corresponding to coupled ORN pairs under the null hypothesis that the coefficients are sampled independently from distinct distributions of positive- and negative-valence uPNs. The null distribution was computed as the product of the PC1 coefficient distributions of positive and negative uPNs (‘marginals’). To take into account the fact that some uPNs participate in multiple coupled pairs (Figure S7), the marginals are calculated by weighing each uPN by the number of pairs it participates in. The resulting marginal distributions for positive and negative uPNs were best fit by Gamma distributions with *κ*_+_ = 1.20, *θ*_+_ = 0.09 and *κ*_−_ = 1.21, *θ*_−_ = 0.05 respectively (Figure 3D).

Next, we calculated the distribution of PC1 readout-angles under the null hypothesis. If the PC1 coefficients of positive- and negative-valence uPNs (*X*_±_) are independently distributed Gamma variables *X*_±_ ∼ Γ(*κ*_±_, *θ*_±_), then their ratio *Y* ≡ *X*_−_*/X*_+_ ∼ *β*^′^ (*κ*_−_, *κ*_+_, 1, *θ*_−_*/θ*_+_), where *β*^′^ denotes the generalized beta prime distribution. The probability density function is,

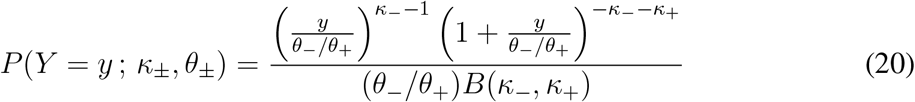

where *B*(·) is the beta function. The PC1 readout-angle for a coupled pair is defined as Φ = tan^−1^*Y* . Transforming variables, we find the null distribution of PC1 readout-angles to be

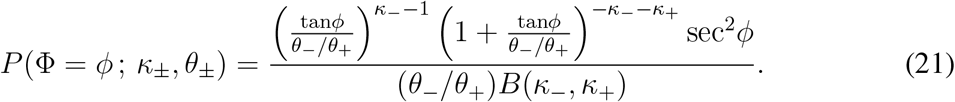

Eq. (21) was used in Figure 3E.

##### 1.2.5 Analysis of AL-to-LH connectivity at the level of individual neurons

To increase the statistical power of our analysis of PC1 coefficient pairs, we analyzed AL-to-LH connectivity at the level of individual uPNs and LHNs. In this Section, ‘uPNs’ and ‘LHNs’ refer to individual neurons instead of neuronal types. As in Section 1.2.1, we neglected neurons that did not participate in any AL-to-LH connections, resulting in a connectivity matrix with 136 uPNs and 1442 LHNs. The weight of each connection was the number of synapse counts between the corresponding uPN and LHN. Performing PCA on the connectivity matrix showed that the first five PCs explained more variance than expected from shuffles (data not shown). As before, we focused on the leading Principal Component (PC1) and analyzed the participation of each uPN in the most prominent pattern of AL-to-LH connectivity.

We identified 62 uPNs postsynaptic to ORNs with known valence (*n* = 30 positive and *n* = 32 negative). There were a total of 124 uPN-pairs that correspond to coupled, behaviorally antagonistic ORN pairs (Figure S8A). To rule out the null hypothesis that the readout weights of coupled ORNs are statistically independent, we tested for a statistical relationship between the empirical PC1 coefficient pairs using Pearson’s chi-squared test for independence (Figure 3F, Figure S8B). To perform the test, the PC1 coefficient pairs were first partitioned into 2-D bins. Let *O*_*ij*_ denote the counts (i.e., number of PC1 coefficient pairs) in the (*i, j*)-th bin. The expected number of counts in the (*i, j*)-th bin, under the null hypothesis of statistical independence, is given by

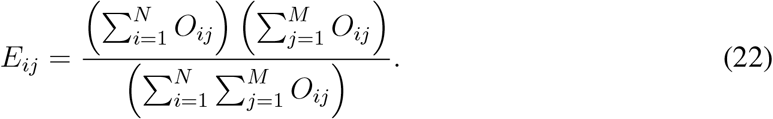

Here *N, M* are the number of bins in the *X* and *Y* directions, corresponding to positive- and negative-valence uPNs, respectively. The value of the *χ*^2^ statistic was calculated as follows,

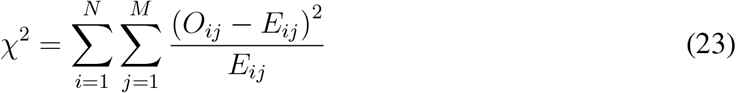

Given the small sample size (*n* = 124), the test statistic does not follow a *χ*^2^ distribution. Thus, we estimated the *P* -values by carrying out the following simulation. First, we generated samples of PC1 coefficient pairs (124 pairs each) by independently sampling from Gamma distributions best fit to the empirical marginals of positive- and negative-valence uPNs (Figure S8A insets; Gamma distribution parameters: *θ*_+_ = 0.046, *κ*_+_ = 0.69, *θ*_−_ = 0.024, *κ*_−_ = 1.40). The generated pairs follow the null distribution (contour lines in Figure S8A). Next, we calculated the *χ*^2^ statistic for the generated samples as described above. Repeating the procedure 10^6^ times, we thus obtained a distribution of the *χ*^2^ statistic under the null hypothesis that the PC1coefficient pairs are statistically independent. The results of the test are summarized in Figure S8B for several bin choices. The empirical *χ*^2^ statistic was significantly greater than that obtained under the null hypothesis, regardless of this choice. This supports the notion that the PC1-coefficient pairs are not statistically independent.

As before, we computed the PC1 readout-angles for coupled ORN pairs (Figure S8C). The null distribution was calculated from the marginals as described in Sec. 1.2.4. We then performed a goodness-of-fit test for the null distribution using the Kolmogorov-Smirnov (KS) test. To circumvent the issue of small sample size, we estimated *P* -values from simulations using the null distribution. We generated samples of PC1 readout-angles (10000 samples, 124 values each) from the null distribution, and computed the KS distance between the null distribution and the sample. The KS distance between the empirical data and the null distribution was significantly larger than that for samples drawn from the null distribution (Figure S8D). This implies that the null distribution does not fully explain the distribution of PC1 readout-angles found in connectomic data.

We note that the distribution of PC1 readout-angles for individual uPNs is skewed towards values larger than *π/*4, contrary to the distribution obtained for uPN-types (Figure 3E). This discrepancy is due to a ‘splitting’ of readout-angles, when the analysis is done at the level of individual neurons (Figure S8E). In summary, this analysis (individual neurons) revealed dependencies between readout-angles downstream from coupled ORNs, while analysis at the level of uPN-types revealed the global bias predicted by our model.

##### 1.2.6 Compound morphometric ratio

The size asymmetry between coupled ORNs is characterized by the following seven morphometric measurements [13]: soma volume and surface area, inner-dendrite volume and surface area, outer-dendrite volume and surface area, and number of dendritic branches. We computed the ratio of the measured values for coupled negative- and positive-valence ORNs [12] for each morphometric quantity 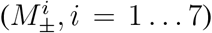. We then calculated the correlation of the resulting morphometric ratios 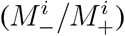 with the PC1 readout-angles of the corresponding uPN pairs [i.e., tan^−1^(*X*_−_*/X*_+_), Figure 3F, Figure S9]. To test the significance of the correlations, we repeated the above procedure for randomly chosen pairs of positive- and negative-valence uPNs. The pairs were generated by randomly pairing uPNs of *opposite valence*, with the constraint that each uPN participates in the same number of pairs as in the original data (repeated 1000 times). We found that inner-dendrite volume, outer-dendrite surface area and dendrite branch number ratios had a significant correlation with readout-angles compared to the controls (*P* values computed as distribution percentiles). In contrast, somatic ratios had a small negative correlation with readout-angles that was statistically significant (Figure S9A,B).

Since the extent of interaction asymmetry likely depends on multiple size asymmetry parameters, we constructed a *compound* morphometric ratio using a linear combination of the ratios of the three dendritic features which were significantly correlated with PC1 readout-angles (Figure 3F). The coefficients in the linear combination were optimized to maximize the correlation between the compound morphometric ratio and PC1 readout-angles. The optimum coefficients for each morphometric ratio are: 0.64 (Inner dendrite volume), 0.15 (Outer dendrite surface area), 0.21 (Dendritic branch number). We found that the correlation between the compound morphometric ratio and PC1 readout-angles is highly significant compared to random uPN pairs.

##### 1.2.7 Analysis of secondary PCs

To check if any additional PCs contribute to valence readout in LH, we computed the difference in the average PC coefficient for positive- and negative-valence uPNs for every PC in the data [*D*(+, −), Figure S10A]. We then calculated the z-score of *D*(+, −) (denoted *z*_*D*_) for each PC by shuffling the empirical connectivity matrix 1000 times. We identified three PCs – PC1, PC4, PC36 – that showed a significant difference between positive- and negative-valence uPNs (|*z*_*D*_| *>* 1.96) relative to both type 1 and type 2 shuffles. However, neither PC4 nor PC36 readout-angles were significantly correlated with *any* morphometric ratios of the corresponding ORNs (Figure S10B,C). Thus, we conclude that valence is primarily readout by the leading Principal Component (PC1) of AL-to-LH projections.

#### 1.3 Model of stimulus classification in AL-to-MB pathway

##### 1.3.1 Modeling a snapshot of AL responses

We defined a ‘snapshot-model’ of ORN responses. Pulse stimuli were initially processed by *N* = 50 ORNs driven by *N* private odorants. ORNs were ephaptically coupled to mimic compartmentalization into sensilla. We then computed the response of each pair of coupled ORNs as follows. Let odor stimuli for one pair of coupled ORNs be denoted by *S*_*A*_, *S*_*B*_, with amplitude 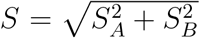 and angle *ϕ*_*S*_ = tan^−1^(*S*_*B*_*/S*_*A*_). The amplitude (*R*) and angle (*ϕ*_*r*_) of the transformed response were computed as,

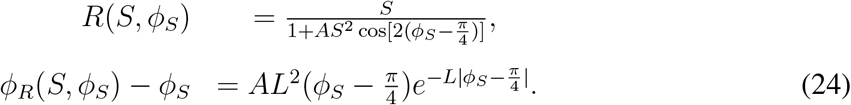

Here, the parameters *A, L* denote the amplitude and range (in stimulus space) of the ephaptic interactions. An illustration of the transformation from *S*_*A*_, *S*_*B*_ to *R*_*A*_, *R*_*B*_ is shown in Figure 4B. The responses *R*_*A*_, *R*_*B*_ can be thought of as the ORN firing rates *x*_*A*_, *x*_*B*_ some time *t* after stimulation. We elected to not use our dynamic model here because parameterizing the interactions using *A, L* shows the effect of ephaptic inhibition on stimulus classification more clearly, compared with the parameters of the dynamic model *K, n* [see Eq. (1) in main text]. Nevertheless we show that the two models impose very similar transformations on the stimulus (Figure 4C, Figure S11A). We neglect local interactions in the AL, and assume that *R*_*A,B*_ represent the activation levels of the corresponding glomeruli.

##### 1.3.2 Assigning labels to stimuli based on their primacy-set

AL responses in our model were not clustered into groups with different labels, as in [18]. Instead we assigned labels to each *N* dimensional stimulus based on its primacy-set, composed of the *N/*2 most active glomeruli. Previous work has shown that the primacy-code is most informative when the size of the primacy-set equals half the dimension of the stimulus space [39]. Glomeruli corresponding to coupled ORNs were assigned opposite valence. The label of a stimulus was set to +1*/*−1 if the majority of glomeruli in the primacy-set had positive/negative-valence, respectively. We also carried out simulations where the stimulus label was not based on a valence majority in the primacy-set, but rather on a random (predetermined) split of glomeruli into two equal groups (Figure S12, triangles vs. squares).

##### 1.3.3 Comparison of clustering based on Bhattacharya distance

We compared the structure of AL responses transformed by ephaptic coupling, to that of responses obtained under the assumption of clustering [18]. For both cases, we computed Euclidean distances (in *N* = 50 dimensional response space) between stimuli with the same label, and stimuli with different labels. In the case of clustered AL responses, the distributions of the two distances were significantly different (Figure 4A). This is because when responses are clustered, nearby stimuli always have the same label. In contrast, the two distance-distributions are very similar for AL responses generated by our model (Figure 4C). This is quantified by the Bhattacharya distance between the distributions (Figure S11B,C).

##### 1.3.4 Random expansion and sparsification simulations

We used the snapshot model to generate responses to sets of *P* = 2500 *N* -dimensional stimuli. We then evaluated the classification performance based on stimuli labels using a Support Vector Machine classifier. At the level of the AL, we repeated these simulations 100 times for each combination of the parameters *A, L*. We compared these results with classification performance in the MB. We computed responses of *N*_*M*_ = 2000 Kenyon Cells (KCs) in the MB as follows. For each simulation, we considered a *N*_*M*_ × *N* dimensional random matrix, which represents random expansion from the AL to MB. Each element in the matrix is sampled i.i.d. from a standard normal distribution. The KC responses to each stimulus were then generated by multiplying the *N* dimensional vector of AL response with the random matrix. The sparsity of KC responses was determined by the parameter *f* . We modeled KC responses to be binary [18], such that a neuron was active if it was among the *f N*_*M*_ neurons with the strongest input. We varied the sparsity level *f* systematically and found that *f* ≈ 0.2 led to optimal classification performance. We repeated the simulations 100 times for each parameter combination (*A, L, f*).

#### 1.4 Initiation of specific behavioral responses

By focusing on the *linear* and *feed-forward* transformation of signals in the AL-to-LH pathway, we have identified alignment of peripheral and central processing that together evaluate valence of arbitrary odor mixtures. Since this alignment was discovered by examining the leading PC of AL-to-LH connectivity, we believe it to be a crucial component of initiation of innate odor-guided behaviors. However, different ‘positive’ odor mixtures may be associated with different behavioral responses [e.g., attraction to food sources, egg-laying, and courtship that were studied in [12]]. We speculate that secondary PCs (which also explain more of the variance of the AL-to-LH connectivity than expected from shuffles, Figure 3A, Figure S6), broad sensitivity profiles of ORNs, nonlinear and recurrent processing could implement the computations necessary for determining what stimulus-specific behavioral responses to produce. More modeling and experimental work is required to understand the such specialized responses, which could additionally be context-dependent.

#### 1.5 Partly structured and partly random AL-to-MB connections

We note that in Ref. [18], a significant boost to MB classification performance was achieved when the AL-to-MB projection matrix was partly structured and partly random. The structured component of that matrix depends on the cluster structure of the inputs, which is absent in our model. Future work will focus on identifying analogous partly structured matrices that can leverage the modified stimulus statistics imposed by ephaptic coupling to further improve performance. We speculate that such a structure could be derived by identifying surfaces in stimulus space where stimulus labels change.

#### 1.6 The role of lateral inhibition in the antennal lobe

Slow lateral inhibition was shown to modulate PN responses in the AL [45]. To keep the dynamic and snapshot models, and the connectome analysis tractable, AL lateral inhibition is not included explicitly in our work. We nevertheless believe that the slow and broadly tuned lateral inhibition in the AL [45] (but not *global*, see [46]) plays two important roles in the context of our modeling work. First, this inhibition could implement the normalization necessary to achieve primacy-coding in the AL, which we use to assign labels to stimuli in Figure 4. Second, we note that to compute the net-valence, signals from positive- and negative-valence ORNs should have readout weights with opposite sign [Eq. (2) in main text]. Broad inhibition could mediate a shift of the baseline value of the readouts. Given the bias in readout weights, an upward shift of baseline would assign the ‘correct’ sign (matching the valence of the ORN) to the readouts.

## 2 Supplementary Figures

**Figure S1.**
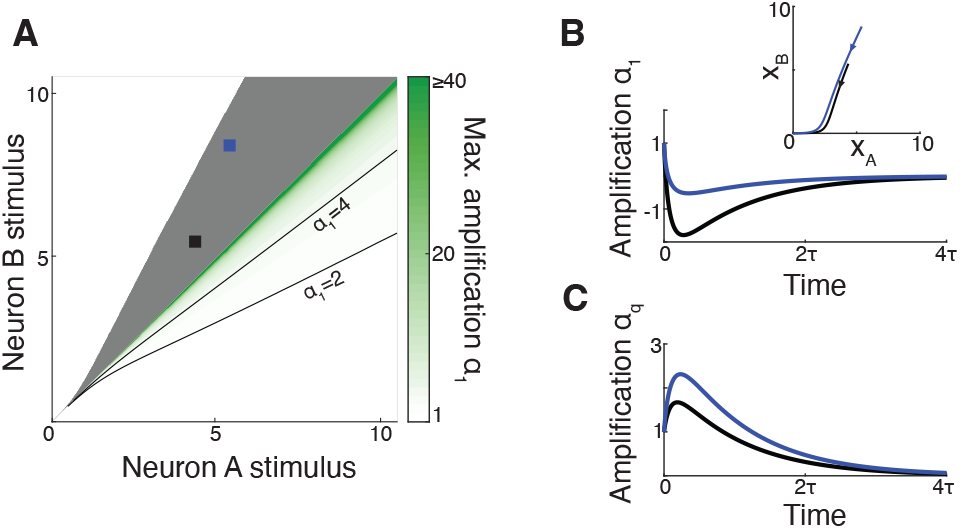
A specific linear combination of paired-ORNs’ firing-rates is necessary for amplification. (A) Maximum valence amplification computed by setting the coefficient *q*^1*/n*^ → 1 [i.e., using 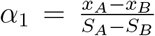, regardless of *q* and *n*]. Not using the sensillum-specific coefficient results in *negative* values of *α*_1_ (gray shaded region), implying that readouts with misaligned weights can lead to incorrect identification of stimulus valence from ORN responses. (B) Temporal evolution of *α*_1_ for two example stimuli, indicated by black and blue squares in panel (A). In both cases, *α*_1_ becomes negative. Inset, firing-rate trajectories resulting from these stimuli. (C) Using the appropriate coefficient in the computation of transient amplification leads to *α*_*q*_ *>* 1 for the same stimuli. Parameters used: *q* = 0.3, *n* = 2, *K* = 1.

**Figure S2.**
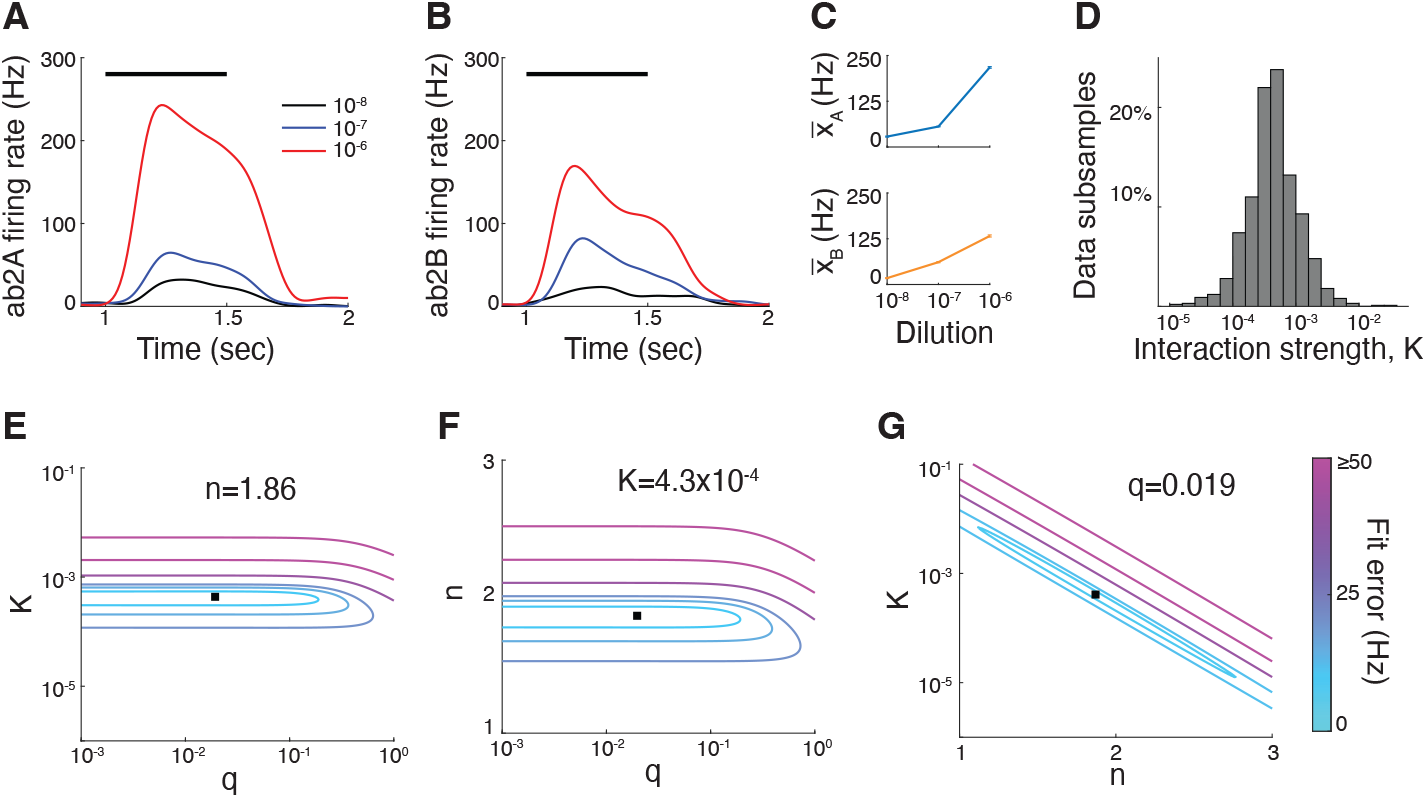
Details of fitting model parameters to ab2 sensillum. (A) ab2A firing rate (averaged over *n* = 9 trials) during a 0.5 s presentation of a single odorant (Methyl acetate, indicated by black bar) at three concentrations (colors). (B) Same as (A), for ab2B. Here the odor presented is Ethyl 3-hydroxy butyrate. (C) Steady-state firing rates 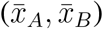 calculated from (A,B) for different odorant concentrations. These steady-state firing rates were used to approximate the respective stimuli strength (*S*_*A*_,*S*_*B*_) for odor mixture experiments (Figure 2E). (D) Histogram of the best fit interaction strength parameter *K* for ab2 odor mixture experiments (1000 data subsamples). (E,F,G) 2D ‘slices’ of the fitting error as a function of the three model parameters, *q, n* and *K*. In each slice, the constant parameter is set to the best-fitting value, indicated in each panel.

**Figure S3.**
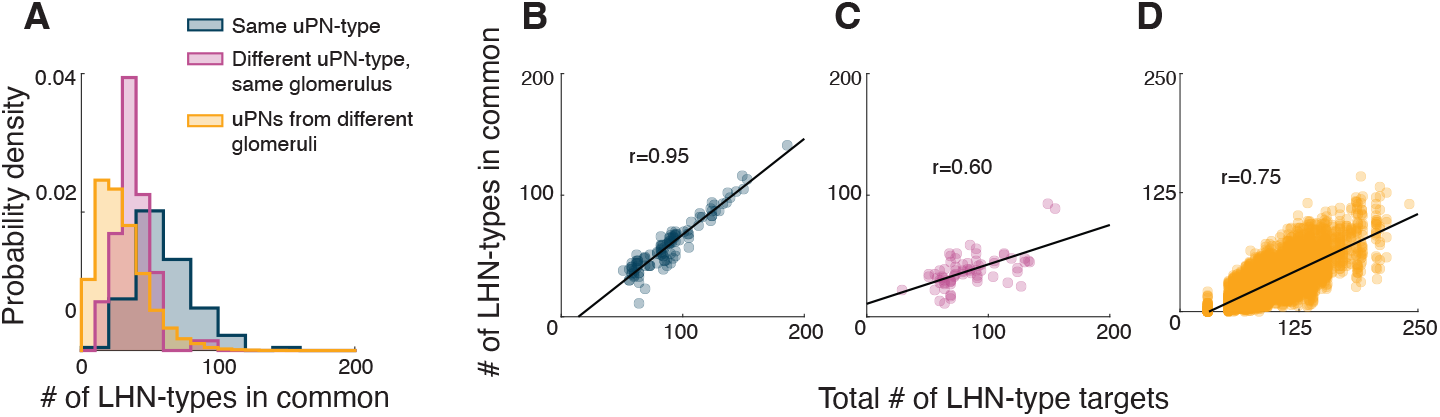
Different uPN-types from the same glomerulus have different LH connectivity. (A) Histogram of the number of target LHN-types in common for two uPNs of the same type (blue), uPNs of different types but originating in the same glomerulus (pink), and uPNs originating in different glomeruli (yellow). Different uPN-types from the same glomerulus have significantly fewer targets in common compared to uPNs of the same type. (B-D) Scatter plots of the number of target LHN-types in common versus (B) the total number of LHN-targets for pairs of individual uPNs of the same type, (C) of different types but from the same glomerulus, and (D) from different glomeruli. Here, for each pair, the total number of LHN-targets was calculated as the minimum of the total number of LHN-types postsynaptic to each uPN in the pair. Also shown are correlation values (*r*) between the two quantities. The number of shared targets is highly correlated with the total number of targets for uPNs of the same type. On average, uPNs of the same type share 79% of targets. In contrast, uPNs of different types from the same glomerulus share only 32% of their targets, while uPNs from different glomeruli share 47% of LHN targets on average.

**Figure S4.**
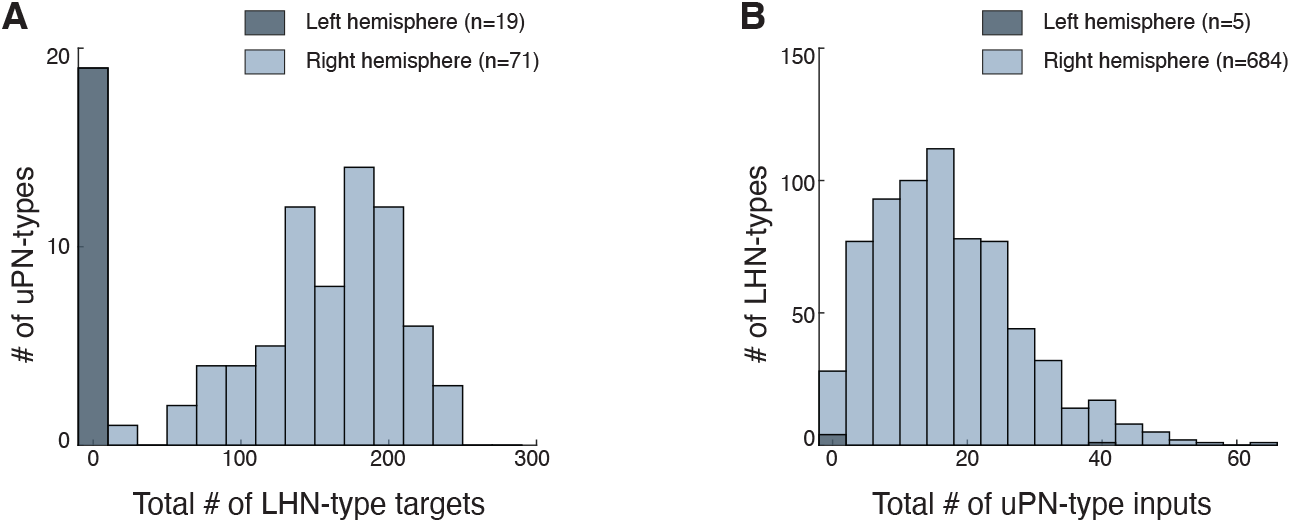
Distributions of the number of outgoing/incoming connections are preserved in the shuffles. Histograms of (A) number of outgoing connections per uPN-type, and (B) number of incoming connections per LHN-type in the empirical AL-to-LH connectivity matrix. Colors differentiate neurons from the left and right hemisphere of the fly brain. Neurons that participate in zero connections were not included in further analyses. Shuffled connectivity matrices were generated under the constraint of preserving both incoming and outgoing connection frequency statistics from the empirical data.

**Figure S5.**
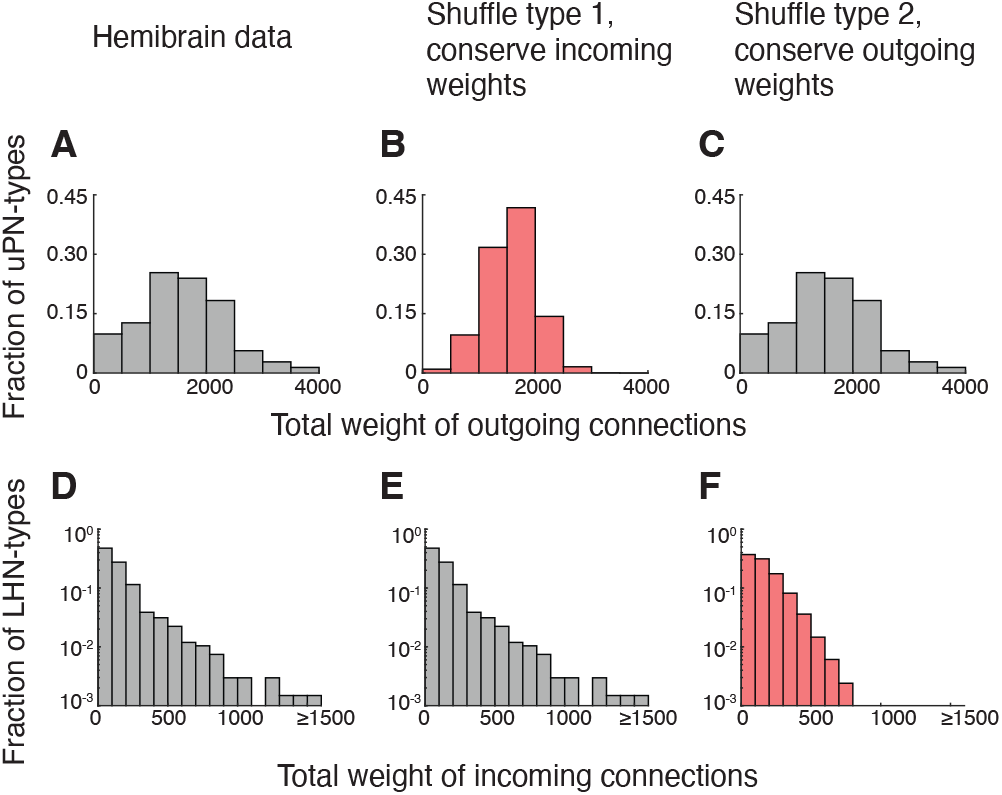
Distributions of outgoing/incoming connection weights are separately preserved in type-1,2 shuffles. (A-C) Histogram of total weight of outgoing connections per uPN in the original dataset, type-1 shuffled datasets and type-2 shuffled datasets respectively (Sec. 1.2.2). Type-2 shuffles preserve the statistics of outgoing connection weights. (D-F) Histogram of total weight of incoming connections per LHN in the original dataset, type-1 shuffled datasets and type-2 shuffled datasets respectively. Type-1 shuffles preserve the statistics of incoming connection weights. Note the lack of outliers (i.e., LHNs that receive strong inputs) for type-2 shuffles (F). Statistics in (B,C,E,F) were generated using 1000 shuffles each.

**Figure S6.**
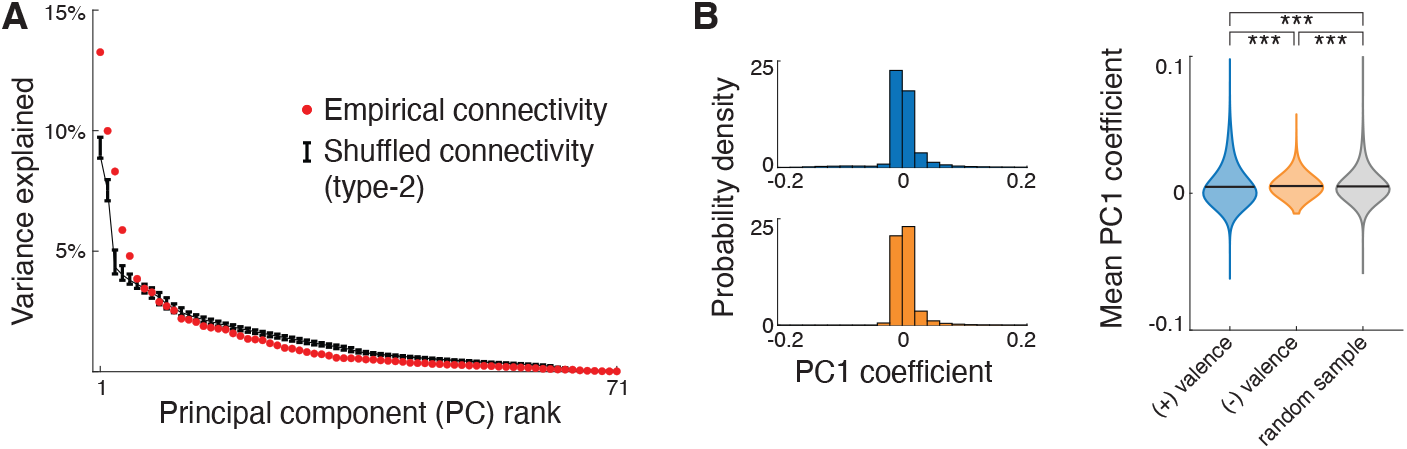
Readout weights of negative-valence uPNs are not biased towards smaller values in shuffled connectivity matrices. (A) Variance explained by Principal Components (PCs) of the empirical AL-to-LH connectivity matrix (red) and type-2 shuffled connectivity matrices that preserve outgoing connection weights (black, error bars indicate 95% confidence intervals from 1000 shuffles). The first 5 PCs of the data explain significantly more variance than those of type-2 shuffles, implying that they represent structured patterns of AL to LH connectivity. (Left) PC1 coefficients of positive- and negative-valence uPNs obtained from type-2 shuffles. (Right) Distribution of the mean PC1 coefficient for positive- and negative-valence uPNs obtained by subsampling uPNs (blue and orange, respectively; 10^4^ shuffled datasets, each subsampled 100 times). Also shown are mean PC1 coefficients for a random sample of 14 uPNs (gray; 10^4^ shuffled datasets, 100 random samples each). Black lines indicate distribution mean. The mean PC1 coefficient of negative uPNs is slightly *larger* than that of positive uPNs and random samples, in contrast to the empirical data (Figure 3B). This difference in the mean PC1 coefficients, though statistically significant, is roughly two orders of magnitude smaller than that seen for the empirical data. ^***^*P <* 0.001, one-sided Student’s t-test. These results are consistent with those obtained for type-1 shuffles (Figure 3A,C).

**Figure S7.**
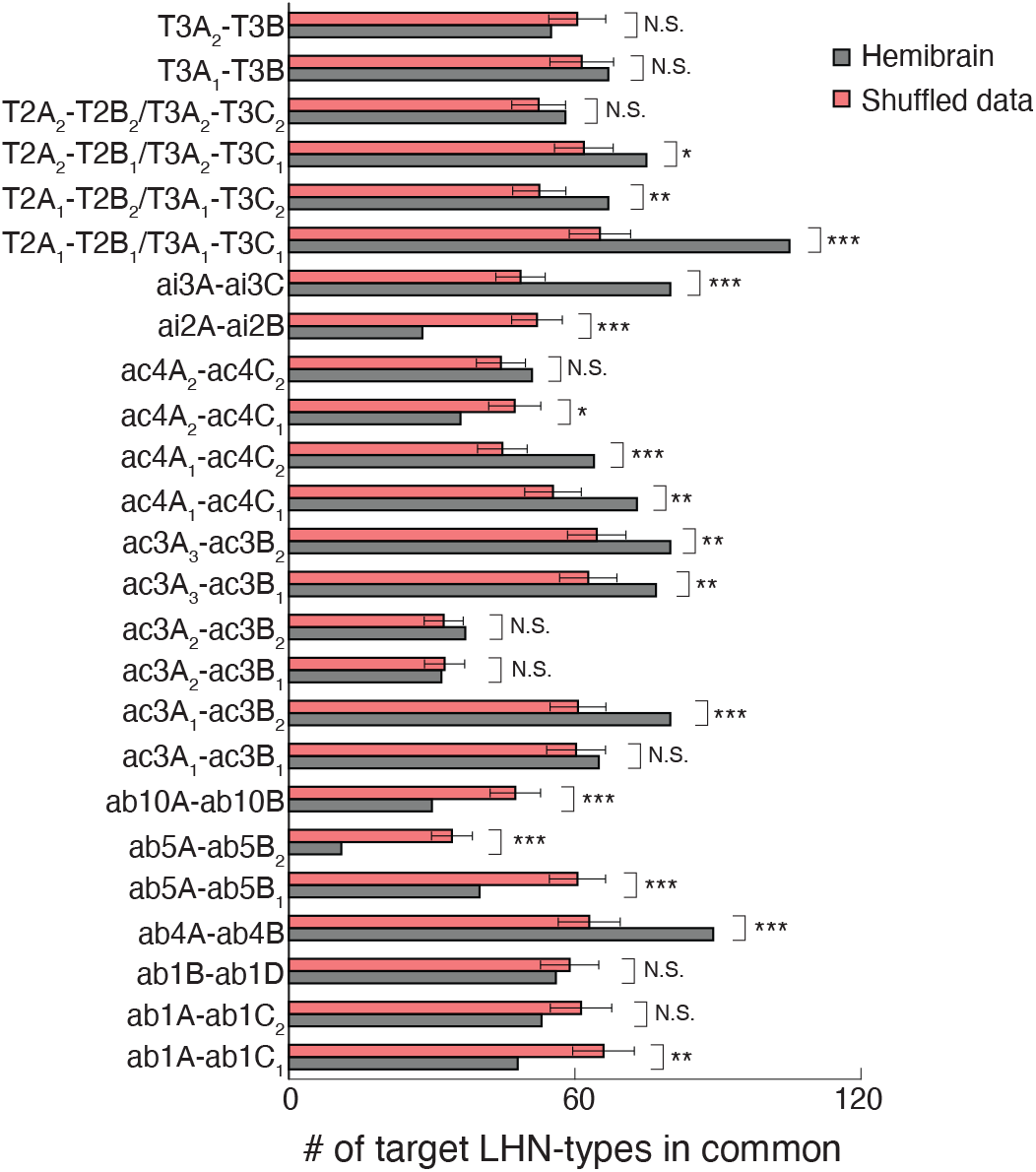
The number of shared LHN targets for uPNs postsynaptic to coupled ORN pairs are significantly different in shuffled datasets. Number of LHN targets in common for each uPN pair (vertical axis) in the Hemibrain data (gray) and shuffled datasets (pink). For 16/25 pairs, the empirical number of shared LHN-targets is significantly different than that expected from shuffled matrices. (^*^*P <* 0.05, ^**^*P <* 0.01, ^***^*P <* 0.001). Subscripts enumerate multiple uPNs per ORN-type (Table 1). T2 and T3 sensilla have two ORN types in common: Glomerulus VA1v (Or47b) corresponds to both T2A and T3A, while glomerulus VA1d (Or88a) corresponds to both T2B and T3C. Thus, uPNs innervating the shared glomeruli are attributed to both ORN types.

**Figure S8.**
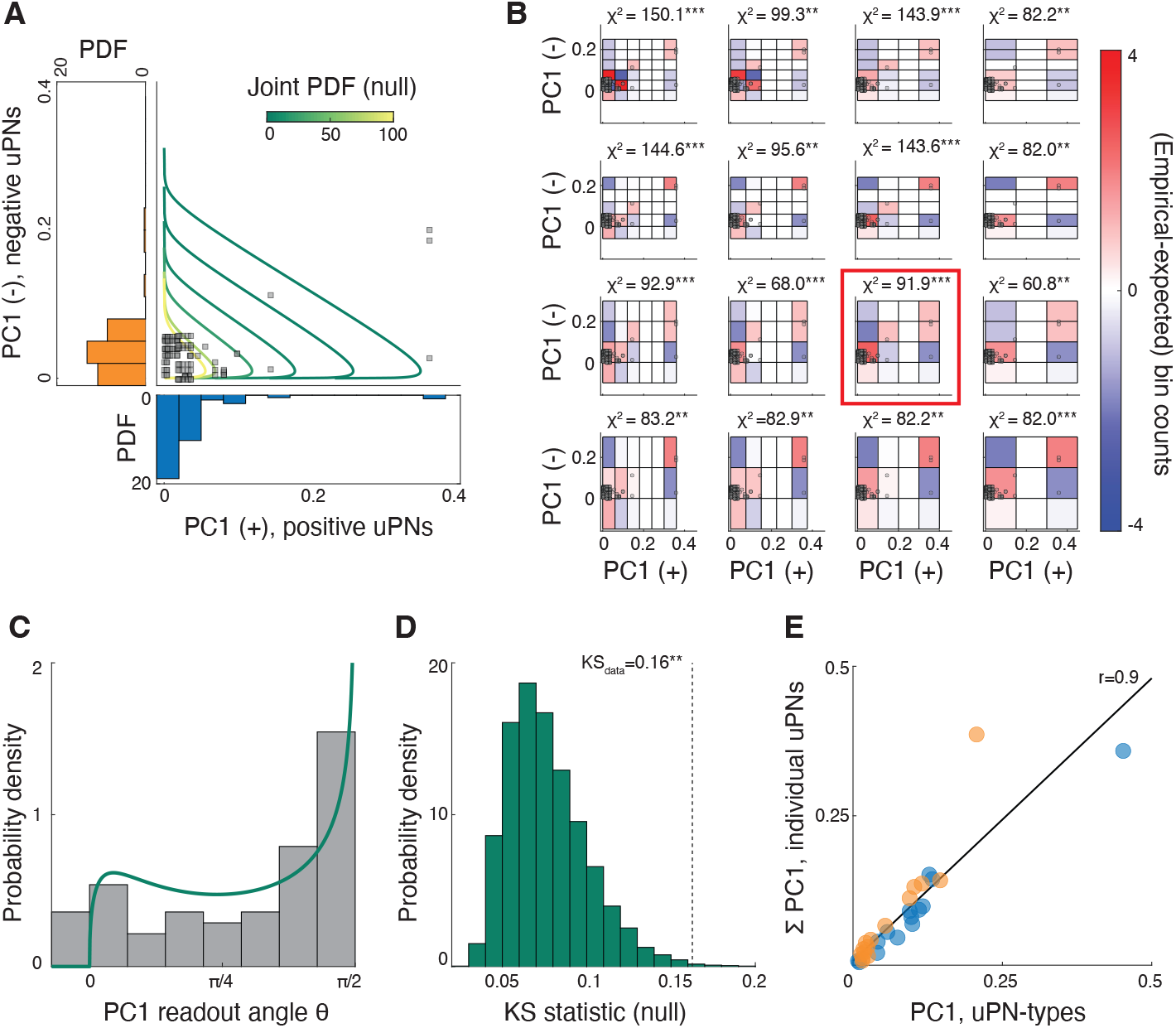
Analysis of AL-to-LH connectivity at the level of *individual* neurons reveals signatures of peripheral interactions. (A) PC1 coefficients of pairs of uPNs postsynaptic to coupled, behaviorally antagonistic ORNs (squares, *n* = 124). Histograms show the marginal distributions of the PC1 coefficients, constructed by counting uPNs once for each pair they participate in. Also shown are contour lines of the null distribution, computed as a product of the Gamma distributions best fit to the marginals. (B) Heat-map of the difference in empirical and expected bin counts (*O*_*ij*_ −*E*_*ij*_, see Sec. 1.2.5) for bin widths. Gray circles: PC1 coefficients for the individual uPN pairs. The empirical *χ*^2^ statistic obtained for each bin choice is indicated (\*\**P <* 0.01, \*\*\**P <* 0.001, *P* -values from simulation with 10^6^ samples). The null hypothesis is consistently rejected. Red box indicates the bin choice in Figure 3 (D) Empirical (gray) and null (green) distribution of PC1 readout-angles. (E) Distribution of KS distance between the null distribution of PC1 readout-angles and samples of readout-angles generated from it (10^4^ samples). The KS distance for empirical PC1 readout-angles is significantly larger, indicating a poor fit with the null distribution. ** *P <* 0.01 (F) Scatter plot of the sum of PC1 coefficients of all uPNs of a given type versus the PC1 coefficient of that uPN-type obtained from the analysis of the neuron-type level dataset. The high degree of correlation (*r* = 0.9) suggests that the connectivity pattern is consistent with the global biases in readouts obtained for the neurontype level dataset. Colors indicate positiveand negativevalence uPN-types.

**Figure S9.**
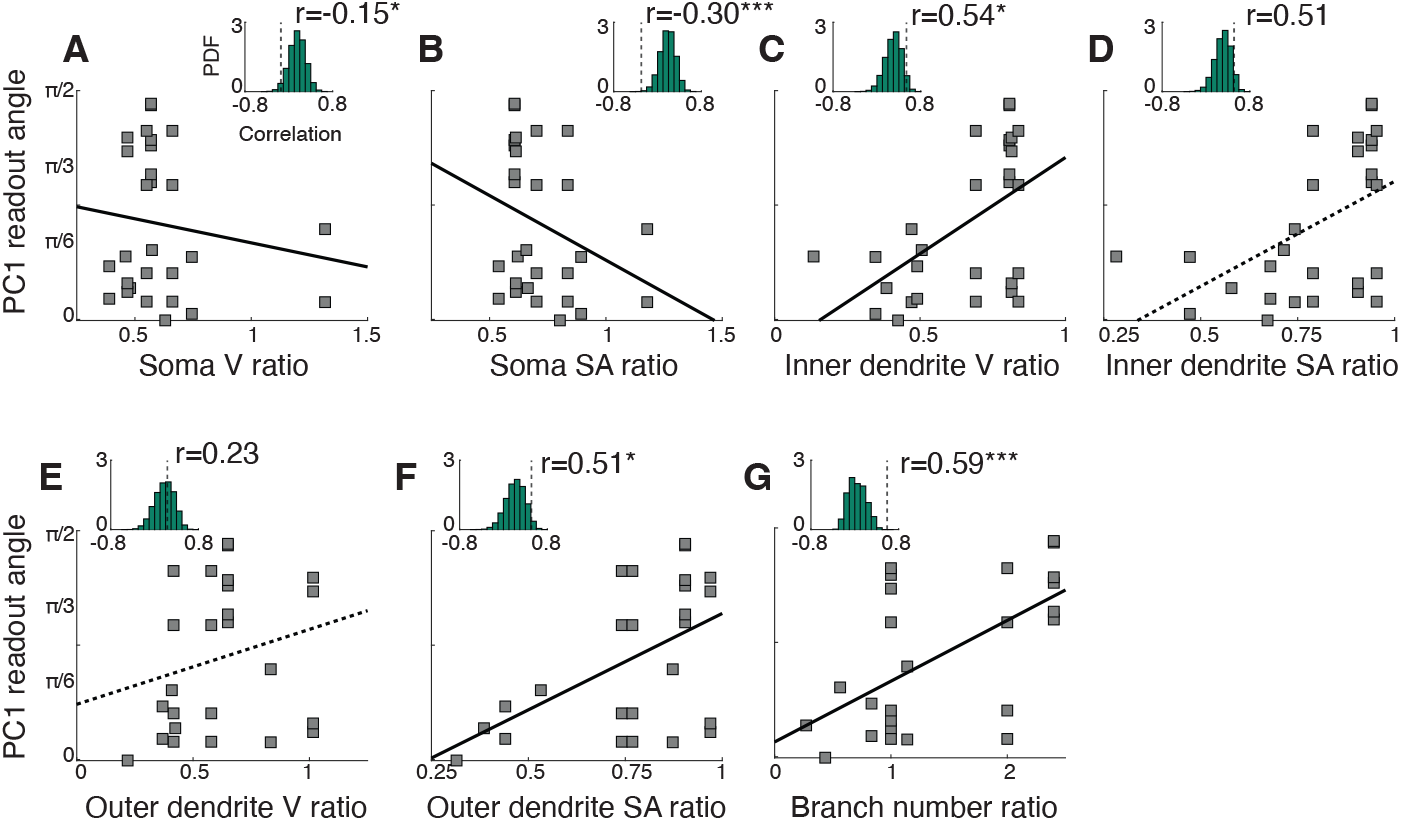
Dendritic, but not somatic, morphometric ratios for coupled ORNs are positively correlated with PC1 readout-angles. Correlation of the PC1 readout-angle with ratios of seven morphometric measurements of the corresponding coupled ORNs, (A) Soma volume; (B) Soma surface area; (C) Inner dendrite volume; (D) Inner dendrite surface area; (E) Outer dendrite volume; (F) Outer dendrite surface area; (G) Number of dendritic branching points. Insets show the correlation between the same morphometric ratio for random, valence opponent ORN pairs and the PC1 readout-angles of their postsynaptic uPNs (10^5^ iterations). We found that inner dendrite volume, outer dendrite surface area, and branch number ratios have a significant positive correlation with PC1 readout-angles.^*^*P <* 0.05, ^***^*P <* 0.001, *P* -values computed from distribution percentiles.

**Figure S10.**
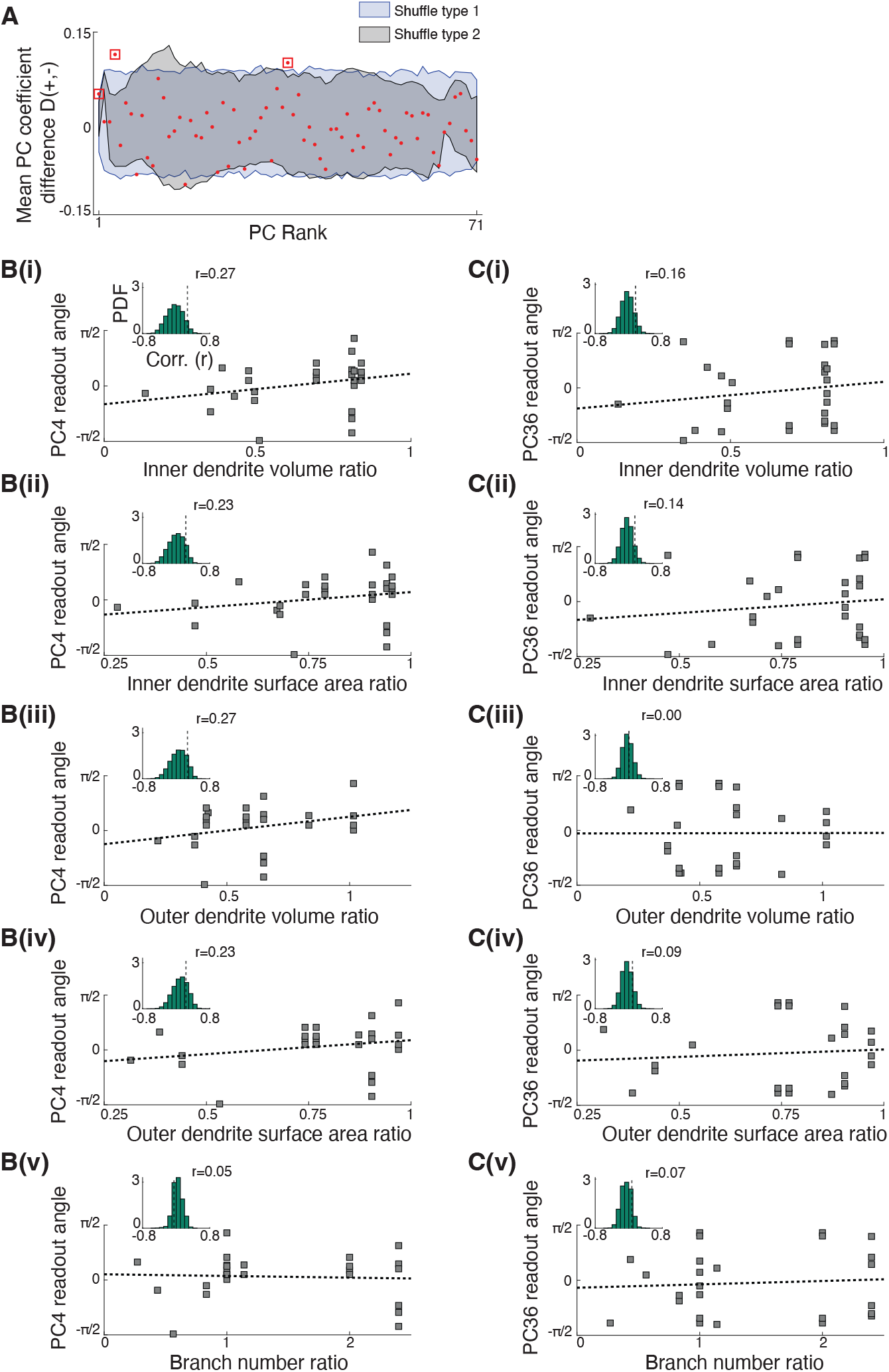
Secondary PCs of AL-to-LH connectivity do not reflect peripheral asymmetries. (A) The difference between mean PC coefficients of positive- and negative-valence uPNs [*D*(+, −), Sec. 1.2.7] as a function of PC rank for the empirical connectivity matrix (red). Shaded regions are 95% confidence intervals of *D* from 1000 shuffled datasets. Aside from PC1, two additional PCs showed a significant difference between positiveand negativevalence uPNs: PC4 and PC36 (red squares). (B) Correlation of dendritic morphometric ratios of coupled ORNs with the PC4 readout-angles of their postsynaptic uPNs. Insets: analogous correlations obtained for random, behaviorally antagonistic ORN pairs. PC4 readout-angles are not significantly correlated with any morphometric ratios (somatic ratios not shown). (C) Same as (B) but for PC36. PC36 readout-angles too are not significantly correlated with any morphometric ratios (somatic ratios not shown).

**Figure S11.**
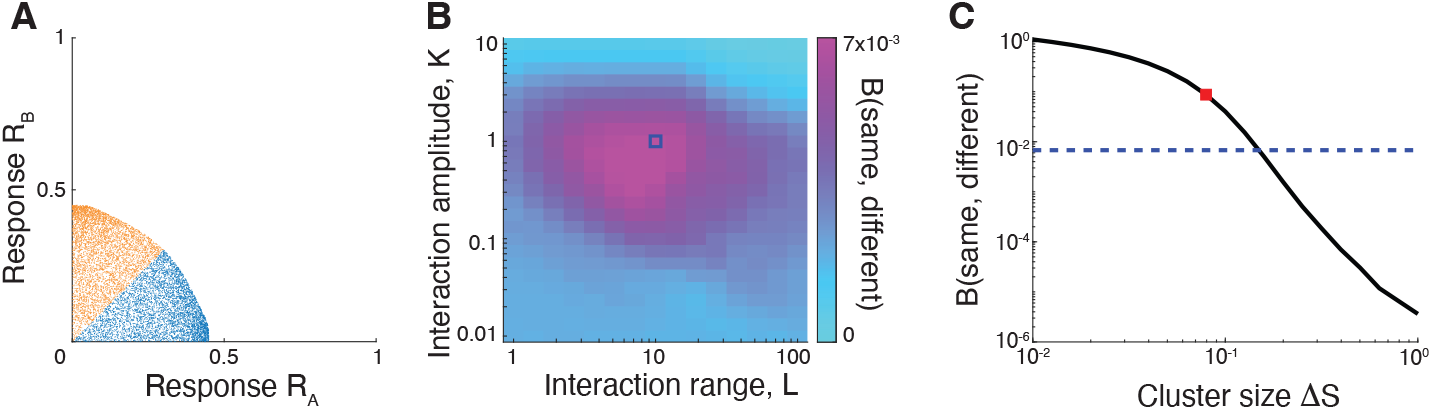
Snapshot model of Antennal Lobe responses. (A) Firing rates of two coupled ORNs from the dynamical model [Eq. (3)] at time *t* = 0.8*τ* . The stimuli here are uniformly distributed, *S*_*A,B*_ ∼ *U* (0, 1). Colors indicate the stimulus label, computed using primacy coding. Here, *q* = 1, *K* = 10, *n* = 6. The distribution of responses is not clustered, and similar to that shown in Figure 4C which was generated using the simpler ‘snapshot’ model [Eq. (24)]. (B) We generated two distributions of distances (in 50D response space) between pairs of responses in the snapshot model: distances between responses to stimuli with the same label; and between responses to stimuli with different labels. Shown here is the Bhattacharyya distance between the two distributions as a function of the interaction parameters in the snapshot model – interaction range (*L*) and amplitude (*A*). A large value of the Bhattacharyya distance implies more dissimilarity between the two distributions. Notably, the region of parameter space with larger Bhattacharyya distances overlaps with the region of high classification performance (blue square, optimal interaction parameters from Figure 4D,E). (C) We generated distributions of distances between pairs of responses having same/different labels under the assumption of AL clustering [18]. Shown here is the Bhattacharyya distance between the distributions as a function of the cluster size (black curve). The blue dashed line corresponds to the Bhattarcharyya distance in the snapshot model for the optimal interaction parameters. Note that a cluster size of Δ*S* = 0.1 [typical value used in [18], red square] corresponds to a Bhattacharyya distance ∼ 30 times larger than that imposed by the snapshot model. This suggests that the structure of AL responses imposed by ephaptic interactions is more subtle than that imposed by AL clustering.

**Figure S12.**
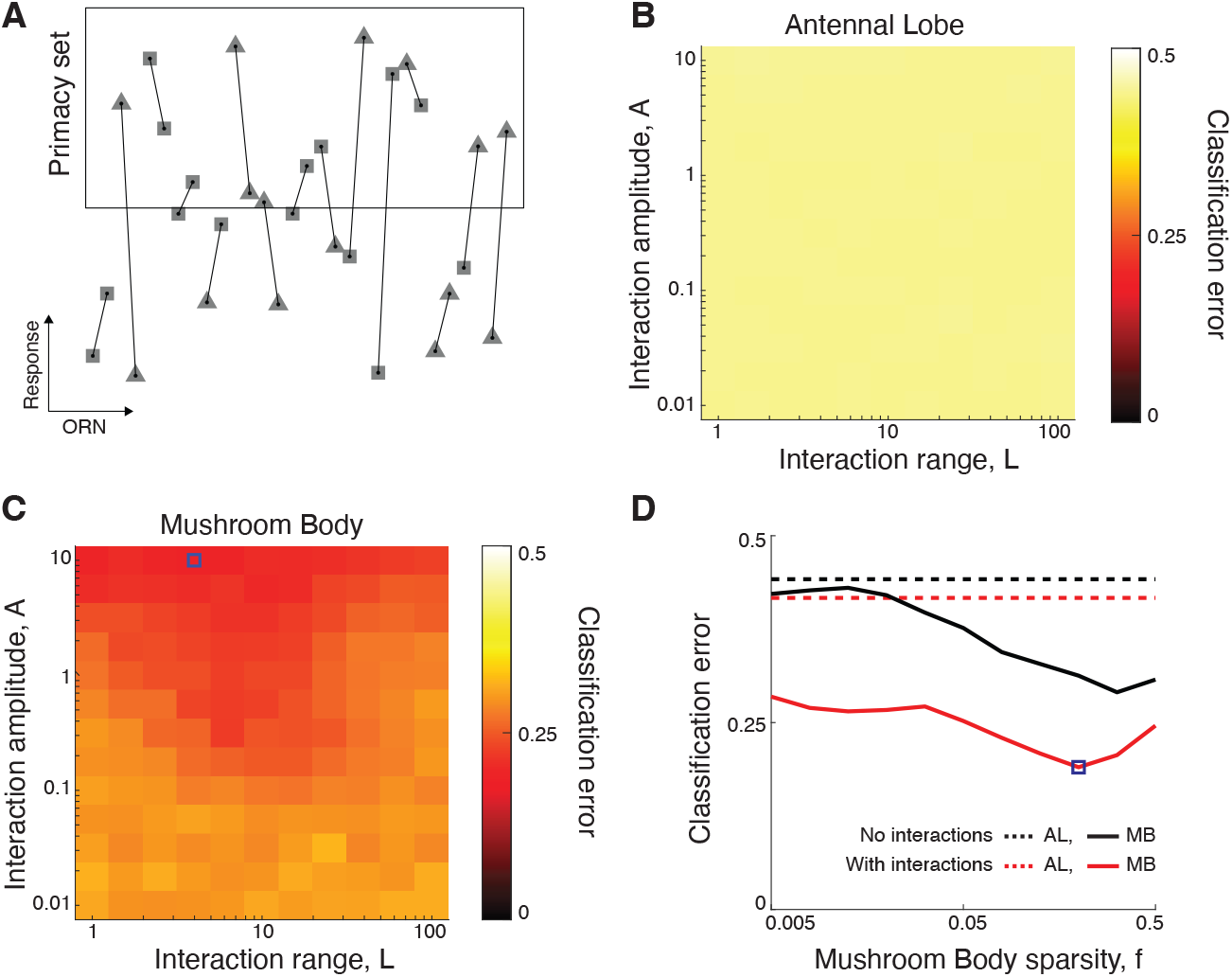
Ephaptic interactions improve classification performance even when labels are based on a random partition of glomeruli within the primacy set. (A) A scenario of the snapshot model where the ORNs are randomly partition into two groups independent of their valence (i.e., triangles and squares). Coupled ORNs do not necessarily belong to opposing groups. Each stimulus is given a label based on whether the primacy-set consists of more triangles (−1) or more squares (+1). (B,C,D) Classification performance in the AL and the MB as a function of the interaction parameters and MB sparsity, as in Figure 4. Ephaptic coupling led to a 2-fold improvement in MB classification in this case, despite the interactions not being ‘aligned’ to the ORN partition. Blue square in (C,D) indicates optimal parameter values.

### 3 Supplementary Table

**Table S1.**
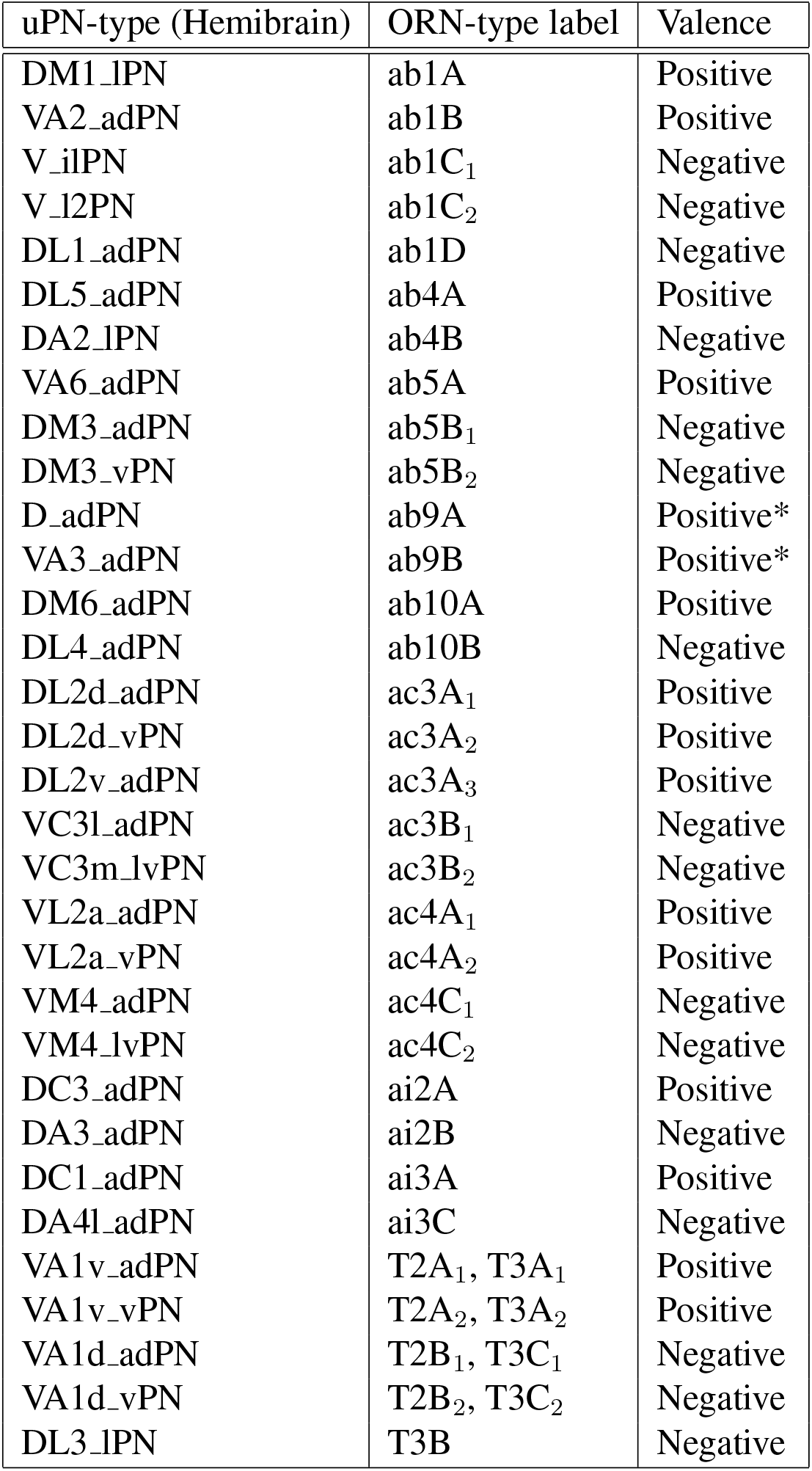
List of uPN-types with known valence used in the analysis. The first column indicates each uniglomerular projection neuron type as labeled in the Hemibrain connectome ([Glomerulus ID] [Cell cluster ID] Projection Neuron, [16]). The second column indicates the corresponding ORN-type [47]. Subscripts 1, 2, … are used to differentiate multiple uPN-types that innervate the same glomerulus. Of note, T2 and T3 sensilla have two glomeruli in common: Or47b/VA1v-projecting ORNs are the A neurons in both T2 and T3 sensilla, while Or88a/VA1d-projecting ORNs are the B neurons in T2 and C neurons in T3. Thus, uPNs innervating these shared glomeruli are attributed to both ORN-types. The third column indicates the valence signal carried by each uPN-type based on their presynaptic ORN valence [Ref. [12] and references therein]. ORNs housed in the same sensillum have opposite valence. A notable exception is the ab9 sensillum, where both ORNs were shown to mediate attraction, i.e., have positive valence [12]. Since the notion of ‘net valence’ is thus unclear for this sensillum, it was not included in our analysis.

